# The biosynthesis of FAD nucleotide analogues in *Escherichia coli* alters its sensitivity to aminoglycoside antibiotics

**DOI:** 10.1101/2025.06.21.660729

**Authors:** Ateek Shah, Prathith Bhargav, Shreya Mahato, Reman Kumar Singh, Yashwant Kumar, Pratishruti Panda, Janhavi Borkar, Lavisha Parab, Nishad Matange, Arnab Mukherjee, Amrita B. Hazra

## Abstract

Cofactors such as flavin adenine dinucleotide (FAD), nicotinamide adenine dinucleotide, coenzyme A, and S-adenosylmethionine cofactors share a common adenosine moiety that is generally not considered to contribute directly to catalysis. In FAD biosynthesis, the adenosine group is installed by the flavin mononucleotide adenylyltransferase (FMNAT) domain of the bifunctional FAD synthetase using FMN and ATP. This study explores the nucleotide selectivity of the *Escherichia coli* FMNAT domain and physiological effects of FAD nucleotide analogues synthesized with cellular nucleotides other than ATP. First, we engineer the FAD synthetase enzyme at an active site loop to produce FAD nucleotide analogues. Chromosomal substitution of the wild-type *E. coli* FAD synthetase with the mutated version results in the intracellular synthesis of these noncanonical FAD nucleotide analogues and confers tolerance to aminoglycoside antibiotics. Upon investigating their potential downstream metabolic roles, we find that their biosynthesis perturbs cellular metabolism and they could bind to a range of *E. coli* enzymes. These findings validate the FMNAT domain loop as a viable site for cofactor engineering, with potential for synthetic biology applications. Finally, the observed correlation between FAD analogue production and increased tolerance to aminoglycosides reveals novel metabolic mechanisms underlying antibiotic sensitivity.

## INTRODUCTION

Flavin adenine dinucleotide (FAD) is an essential redox cofactor across the domains of life. The biosynthesis of FAD originates from the nucleotide metabolism pathway - guanosine triphosphate converts to 5-amino-6-(D-ribitylamino)-uracil via pyrimidine intermediates and ribulose-5-phosphate, forming the pteridine intermediate 6,7-dimethyl-8-ribityllumazine.^1^ In the final step, two molecules of the pteridine intermediate are dismutated by riboflavin synthase, forming the isoalloxazine ring that yields riboflavin (vitamin B_2_) along with 5-amino-6-(D-ribitylamino)-uracil.^1^ Subsequently, riboflavin undergoes phosphorylation by riboflavin kinase to generate flavin mononucleotide (FMN).^2,3^ FMN is then adenylated to form FAD by the action of FMN adenylyltransferase (FMNAT) (Figure 1A).^2^ The redox properties of FAD can be attributed to the isoalloxazine ring, whereas the adenosine part is currently viewed as a binding and recognition handle.^4–6^

**Figure 1.**
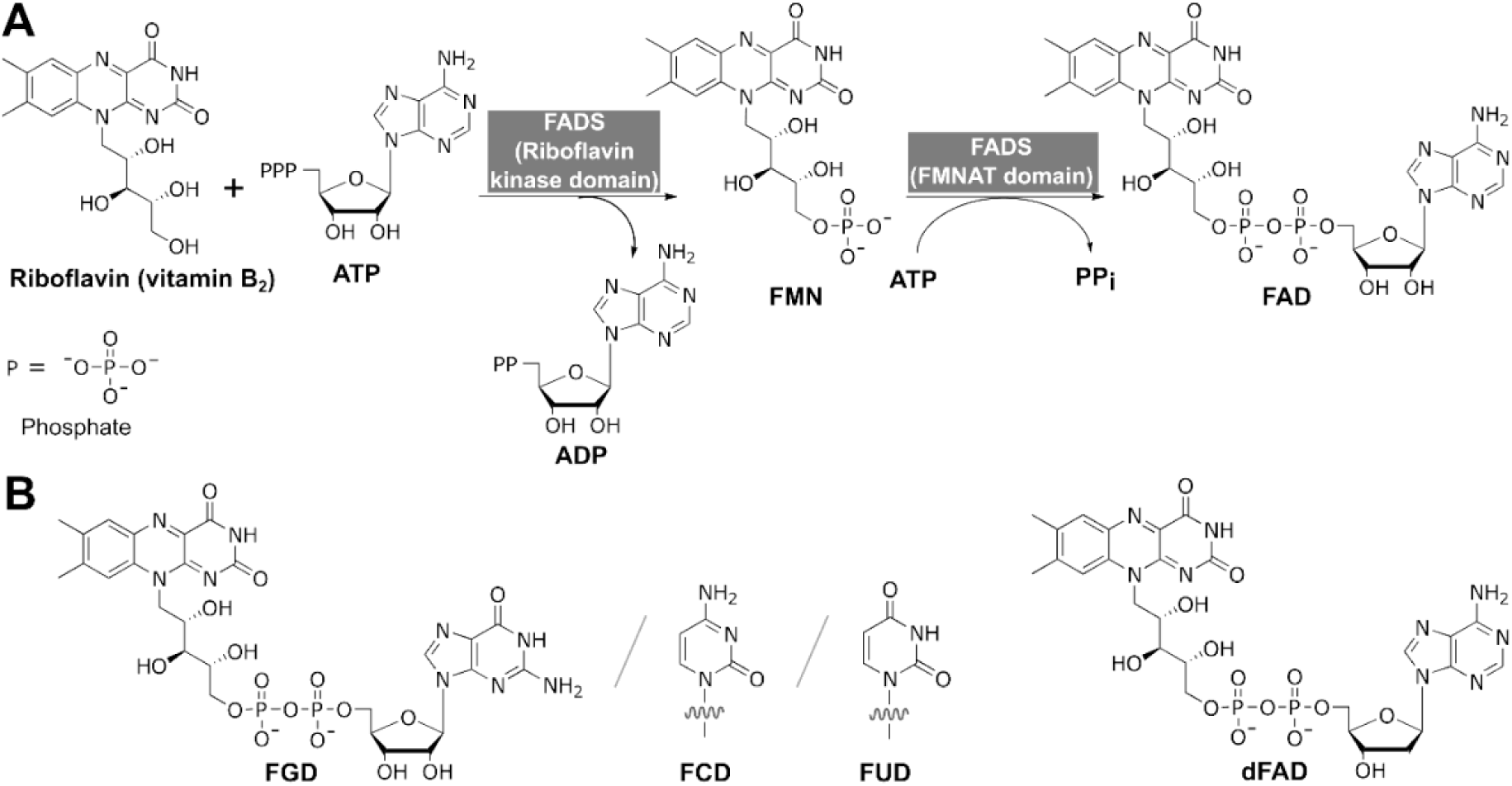
Biosynthesis of flavin adenine dinucleotide (FAD) and its nucleotide analogues. **(A)** The biosynthesis of FAD involves conversion of riboflavin (vitamin B_2_) to flavin mononucleotide (FMN), followed by adenylation to produce flavin adenine dinucleotide (FAD). Both steps use adenosine triphosphate (ATP) as a co-substrate and produce adenosine diphosphate (ADP) and inorganic pyrophosphate (PP_i_) as byproducts. **(B)** FAD nucleotide analogues - flavin guanine dinucleotide (FGD), flavin cytosine dinucleotide (FCD), and flavin uracil dinucleotide (FUD). When adenosine is replaced by deoxyadenosine, deoxy-flavin adenine dinucleotide (dFAD) is formed. Even though these FAD analogues do not naturally occur in cellular systems, the starting substrates FMN and nucleotides are available in cells. Hence, this set of FAD nucleobase/nucleotide analogues can, in principle, be generated in cells using the appropriate FMN adenylyltransferase (FMNAT) enzyme.

The adenosine moiety, also found in other cofactors such as nicotinamide adenine dinucleotide (NAD), S-adenosyl methionine (SAM), and coenzyme A, has been considered as a possible remnant of covalently bound coenzymes in an RNA-centric world.^6,7^ Interestingly, hypoxanthine nucleobase analogues of FAD and NAD have been found in cell extracts of marine bacterium *Desulfobacula toluolica* Tol2, along with acetyl-inosino-CoA and benzoyl-inosino-CoA, analogues of coenzyme A.^8^ In FAD, the adenine is found to be at a distance of ∼11-16Å from the isoalloxazine ring when bound to flavoproteins, and appears to play no direct role in its catalytic function.^9,10^ Hence, replacing the adenine by alternate ligands at the nucleobase position finds applications in fluorescence quenching and crystallography studies.^11–14^ Similarly, novel analogues of SAM and NAD modified at the nucleobase or nucleoside region exist, with applications including imaging and structural probes, and enzyme inhibitors.^15,16^ More recently, the cytosine analogue NCD has been used as a synthetic biology tool for driving reductive carboxylation of pyruvate to L-malate, leading to CO_2_ fixation.^17^ On these lines, FAD nucleobase analogues with guanine, cytosine, uracil and deoxyadenosine, namely FGD, FCD, FUD, and dFAD, are active redox cofactors for glutathione reductase, and can be envisioned as potential non-canonical redox cofactors (NCRC) for modified enzymes within a cell, similar to NCD.^10,18^

Non-canonical FAD analogues demonstrate a range of applications such as biosensing, covalent immobilization of flavoproteins, and enzyme electrodes.^10,13,19–22^ FAD is produced in the cell from riboflavin via two steps that each consume a molecule of ATP (Figure 1A).^2^ While the change of nucleotide substrate for riboflavin kinase in the first step should not affect the product outcome as it is a phosphorylation reaction, the incorporation of other nucleotides (GTP/ CTP/ UTP) by the FMNAT will result in FGD, FCD, or FUD (Figure 1B). Riboflavin kinase and FMNAT typically exist as two distinct enzymes in eukaryotes and are fused to form the bifunctional FAD synthetase in prokaryotes.^3,23^ While most riboflavin kinases have been experimentally tested only with ATP, the other nucleotide triphosphates behave as substrates in some cases. Homologues of *Bacillus subtilis* and *rat intestinal mucosa* riboflavin kinase display the ability to use a range of NTPs to varying extents to make FMN^24,25^, whereas some archaeal homologues exhibit specificity for CTP over ATP^26,27^, and a fungal *Rhizopus javanicus* homologue utilizes GTP specifically, while ATP gives no reaction.^28^ In the case of FMNAT, the *Methanocaldococcus jannaschii* FMNAT shows *in vitro* activity with nucleotides other than ATP resulting in FAD and its nucleobase analogues.^10,29^ As ATP, GTP, CTP, UTP, and the corresponding dNTPs exist naturally in the cell, non-canonical FAD nucleotide analogues can, in principle, be produced in cells if the FMNAT enzyme/domain can accept them as alternate substrates.

The goal of this study was to create an *E. coli* strain that can synthesize nucleotide analogues of FAD *in vivo* using cellular nucleotides. This strain will act as a model system to explore the physiological role of the nucleotide part of FAD, and test for potential applications of FAD analogues as biorthogonal cofactors. For this, first, we altered the FMNAT domain of the *E. coli* FAD synthetase (*Ec*FADS) enzyme *in vitro* to produce FAD analogues and accordingly engineered the enzyme within the *E. coli* chromosome to produce FAD and its nucleobase analogues. We then examined its metabolism with a focus on redox functions and found that the strain exhibits decreased sensitivity towards aminoglycosides such as kanamycin, amikacin, streptomycin, and gentamicin. As redox biochemistry plays a central role in antibiotic resistance, we attribute this altered sensitivity to alterations in metabolism arising from the presence of the FAD analogues. Our findings not only establishes a general method to create a strain that produces cofactor analogues but also reveals an unprecedented link to antibiotic sensitivity, which requires detailed mechanistic inspection.

## RESULTS

### *Ec*FADS shows *in vitro* activity with several nucleotides

*Ec*FADS has been functionally and kinetically characterized for its ability to incorporate riboflavin, FMN and its analogues, and ATP.^30^ In contrast, the *Corynebacterium ammoniagenes* FAD synthetase (*Ca*FADS) has been extensively studied structurally and functionally as a model of prokaryotic FAD synthetases.^31–34^ Sequence alignment of *Ec*FADS and *Ca*FADS shows 31% identity and 50% similarity, and the active sites for riboflavin, FMN, and both ATPs are well conserved (Figure 2A). Since no experimental structure is available for *Ec*FADS, we used homology modelling and obtained a high confidence score structure (Figure 2B).^35,36^

**Figure 2.**
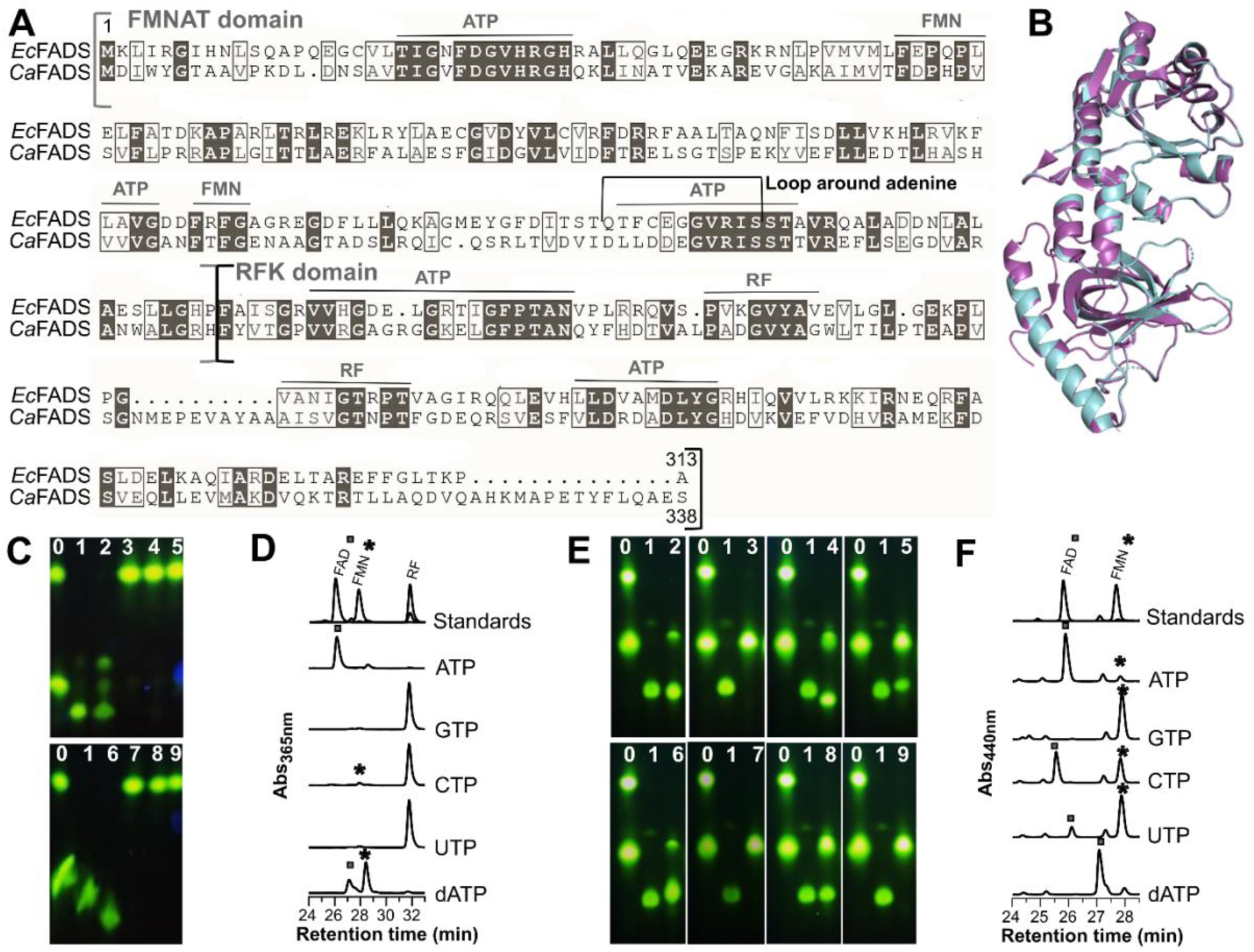
Sequence and modelled structure of *Ec*FADS and its activity with nucleotides. **(A)** Sequence alignment of *Ec*FADS with *Ca*FADS shows 31% sequence similarity and 50% homology. The active site residues (indicated in black boxes)are well-conserved. The square brackets indicate the two domains - FMNAT and riboflavin kinase. The grey horizontal lines indicate the FMN, riboflavin, and ATP-binding regions **(B)** The alignment of modelled *Ec*FADS (shown in cyan) with crystal structure of *Ca*FADS (PDB 2×0k) (shown in magenta) shows complete overlap and a 100% confidence score. **(C)** *Ec*FADS with riboflavin and 3.5 mM nucleotides and deoxyribonucleotide, monitored using fluorescence thin-layer chromatography (TLC).**^‡^ (D)** HPLC analysis of these samples shows activity with ATP, CTP, and dATP (* indicates the FMN and ▪ indicates FAD and its analogues). **(E)** *Ec*FADS with FMN and 3.5 mM nucleotides and deoxyribonucleotide, monitored using fluorescence TLC. Signals for FAD, FCD, FUD, dFAD, and dFCD were observed.**^‡^ (F)** HPLC analysis of these samples shows activity with all nucleotides and dATP except GTP (* indicates the FMN and ▪ indicates FAD and its analogues). **^‡^(Key for C and E)** Fluorescent TLC of *Ec*FADS activity Lane 0- RF and FMN standard, 1- FAD standard, Lanes 2-9, reaction with 2 - ATP, 3 - GTP, 4 - CTP, 5- UTP, 6 - dATP, 7- dGTP, 8- dCTP, 9 – dTTP.

We purified *Ec*FADS and optimized the reaction conditions for riboflavin to FAD conversion with ATP as the co-substrate (Figure S1A, B, C). We then tested a panel of ribonucleotide and deoxyribonucleotide co-substrates ATP, GTP, CTP, UTP, dATP, dGTP, dCTP, and dTTP at 3.5 mM each, which is the cellular concentration of ATP in *E. coli*.^37^

Fluorescence thin-layer chromatography (TLC) and high performance liquid chromatography (HPLC) analysis showed FMN formation with ATP, CTP, and dATP, and FAD and dFAD formation (Figure 2C, D, Figure S1D). Interestingly, when the cellular concentrations of each of the nucleotides were used with riboflavin as a substrate, only dATP continued to show the formation of dFAD (Figure S1E). ^37^

Next, we used FMN as a substrate with the panel of nucleotides to test only the FMNAT activity. We observed that the co-substrates ATP and dATP show high conversion to FAD and dFAD, while CTP, UTP, and dCTP showed partial conversion to FCD, FUD, and dFCD, respectively (Figure 2E, F). Based on these results, as the nucleotides and dATP showed significant conversion in both experiments, we proceeded with these as our co-substrates for subsequent reactions.

### Mutational analysis to alter the nucleotide specificity of the *Ec*FADS FMNAT domain

The *Ec*FADS FMNAT domain interacts with ATP via phosphate, the ribose ring, and the adenine base (Figure S2A, green, yellow and red residues, respectively). To probe the interactions important for ATP specificity and predict mutations, docking followed by molecular dynamics simulation was conducted. S165K was predicted to lower the enzyme’s interaction energy with ATP, GTP and CTP by increasing interaction with the phosphate, while the G23K mutant was predicted to have a decreased preference for ATP and an increased preference for GTP as compared to the wild-type *Ec*FADS (Figure S2B). However, when these mutants were cloned, expressed, purified, and tested, an overall decrease in activity with ATP as compared to the WT *Ec*FADS was observed and no additional promiscuity with nucleotides was seen (Figure S2C). Next, we created the G23S mutant corresponding to a similar mutation G105S in the FtsZ enzyme, which converts it from a GTPase to an ATPase (Figure S2D, the glycine is colored in red).^38^ However, we found it completely inactive (Figure S2E).

### A nucleobase region loop in the *Ec*FADS FMNAT domain plays a role in nucleotide selectivity

A structure-based approach was next employed to identify other nucleotidyltransferases that are homologous to *Ca*FADS and do not utilize ATP. A DALI search with a stringent z-score cutoff of 10 yielded four candidate cytidyltransferase structures (Table S1).^39^ Interestingly, the *E. coli* FMNAT domain and cytidylyltransferases fall within the HxGH motif-containing nucleotidyltransferase superfamily and display high overall structural similarity with only a distinct difference in a loop adjoining the nucleobase (Figure 3A). Glycerol-3-phosphate cytidylyltransferase has specificity for CTP or dCTP, hence, we hypothesized that changing the FMNAT domain loop with that of the cytidyltransferase may result in CTP specificity.^40^ We swapped the *E. coli* FMNAT domain loop (residues 155-166, QTFCEGGVRIS) sequence with the PRTEGIS loop from the glycerol-3-phosphate cytidylyltransferase and purified the resultant *E. coli* Loopswap-FAD synthetase enzyme. On testing its activity with the nucleotide panel and FMN, Loopswap-FAD synthetase displayed similar efficiency at producing FAD as the wild-type *Ec*FADS, and increased ability to utilize GTP, CTP, UTP, and dATP to produce FGD, FCD, FUD, and dFAD, respectively (Figure 3B, S3). Compared to the wild-type FAD synthetase, the Loopswap-FAD synthetase is 17x, 1.2x, and 1.4x as efficient with GTP, CTP, and UTP, respectively (Figure 3B).

**Figure 3.**
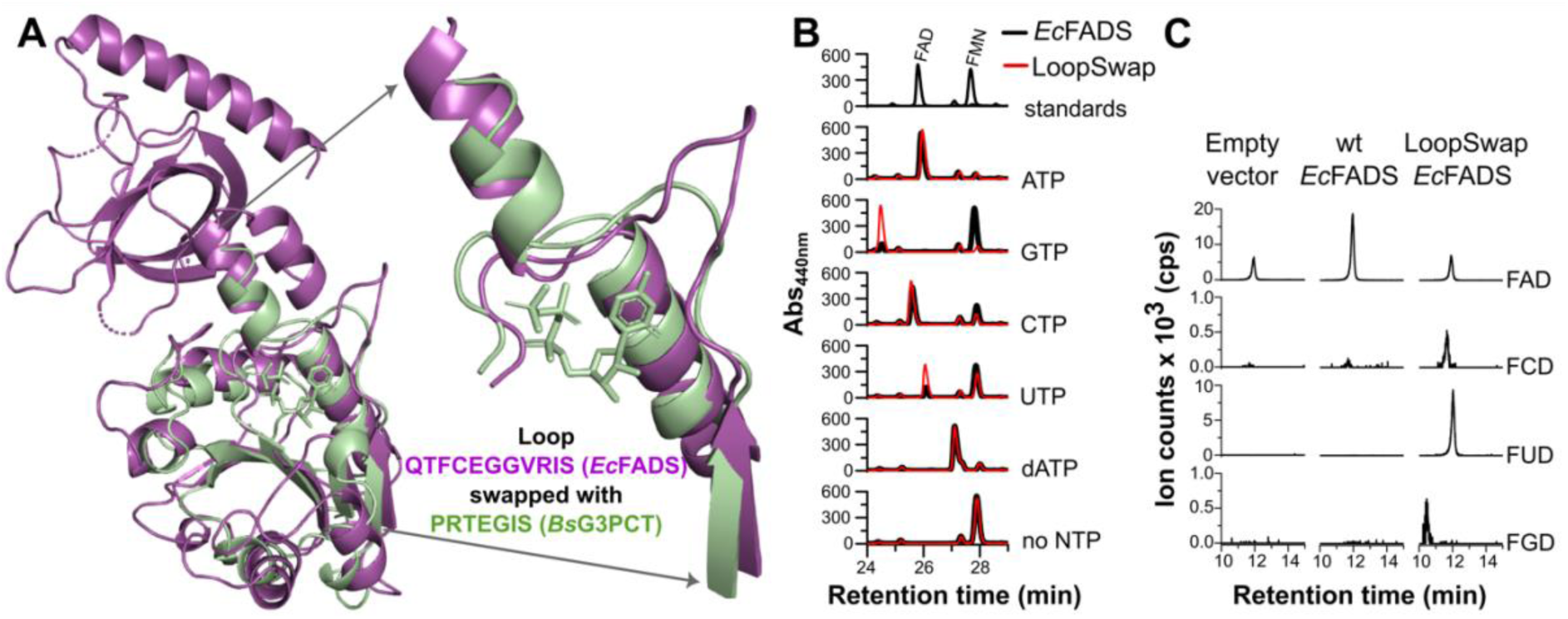
Testing the promiscuity of *Loopswap* mutant *in vitro* and by heterologous expression. (**A**) Structural alignment of *Ec*FADS (magenta) with *Bacillus subtilis* glycerol 3-phosphate cytidylyl transferase (*Bs*G3PCT: green) (PDB 1COZ). The *Ec*FADS FMNAT domain shares high structural similarity with the cytidylyl transferase and very low sequence similarity. The loops surrounding the nucleotide in *Ec*FADS-FMNAT domain and *Bs*G3PCT have different sizes. **(B)** HPLC analysis showing comparative *in vitro* activity of the *Ec*FADS and *E. coli* Loopswap enzymes. The Loopswap enzyme utilizes CTP, UTP and GTP as substrates, while the *Ec*FADS shows lower activity with CTP and UTP and no activity with GTP. (**C**) Extracted ion chromatograms (EIC) of FAD and its analogues from strains heterologously expressing the empty pET28a(+) vector, *Ec*FADS, or the Loopswap enzyme. Expression of the Loopswap enzyme shows signals for FAD nucleobase analogues along with FAD while the one expressing the WT *Ec*FADS shows only FAD.

Next, we heterologously expressed the wild-type and Loopswap-FAD synthetase enzymes in *E. coli* BL21(DE3) and their lysates were analyzed by liquid chromatography-mass spectrometry (LC-MS), with the empty vector as a control. While the lysates of either the empty vector or the wild-type FAD synthetase show only FAD formation, the Loopswap-FAD synthetase lysate additionally showed FCD, FUD, and FGD signals along with FAD (Figure 3C). Thus, our *in vitro* and heterologous expression data indicates that the FMNAT domain loop plays an essential role in nucleotide selectivity.

### The FMNAT loop sequence is important for catalysis and nucleotide specificity

To probe the loop length and the role of the residues in the FMNAT domain loop, we created six truncated loop variants by comparing the sequence, structure and length with the glycerol-3-phosphate cytidylyltransferase loop sequence – QTFCEGIS, QTFCGIS, QTFCIS, QTCGIS, QTFIS, and QTIS (Figure 4A). The QTFCGIS enzyme shows formation of FAD, FUD, and FCD between 20-45% of that of the Loopswap enzyme, whereas the QTFIS and QTIS enzymes showed significant activity with only ATP and dATP (Figure 4B, S4). Overall, for the Shortloop enzyme mutants, a decrease in the formation of FAD and its analogues as compared to the wild-type and Loopswap enzymes was noted (Figure 4B, S4). The decrease in the activity of Shortloop enzymes may be attributed to their low interaction energy with the nucleotides, as calculated for the mutant QTFCGIS with ATP, CTP, and GTP (Figure S2B).

**Figure 4.**
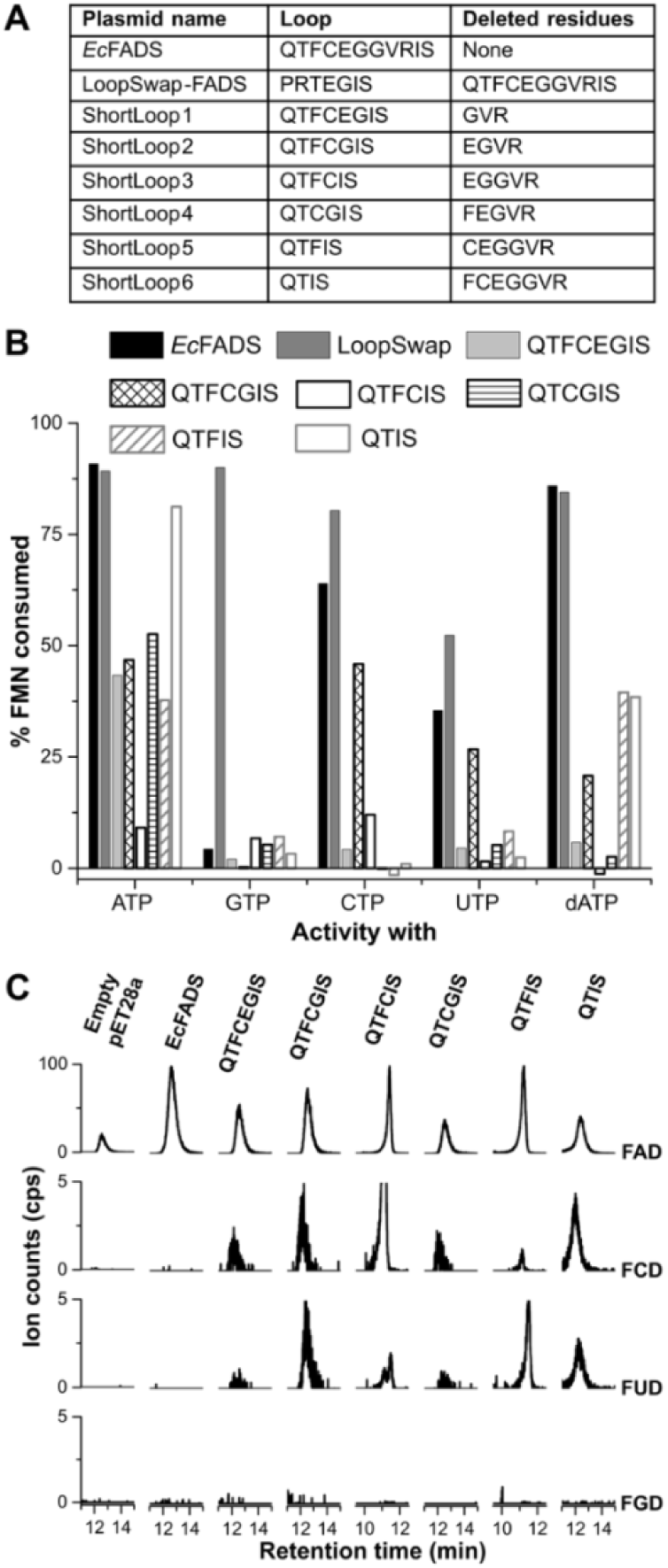
Comparative analysis of the *in vitro* activity of the Loopswap and Shortloop mutants. **(A)** A series of shortened loop sequences were created from the Loopswap. The list of loops, deleted residues, and plasmid names is provided. **(B)** A comparative analysis of FAD and its analogues formed by *Ec*FADS, Loopswap, and Shortloop mutants with various nucleotides, derived from HPLC data. The Loopswap enzyme shows highest activity with other nucleotides as compared to *Ec*FADS. In contrast, the Shortloop mutants show decrease in activity with ATP and other nucleotides. The QTFCGIS mutant is interesting because it shows loss of activity overall but mirrors the Loopswap enzyme’s preference for ATP, CTP, and UTP. **(C)** EIC of FAD and its analogues extracted from the strains heterologously expressing the empty pET28a(+) vector, *Ec*FADS or the Shortloop enzymes. Note that the Shortloop mutants, being less efficient *in vitro*, can yet synthesize FAD nucleobase analogues in the cell when heterologously expressed. This shows that the loop plays a primary role in nucleotide specificity in the cell.

Next, we heterologously expressed the Shortloop enzymes in *E. coli* BL21(DE3) cells to understand their cellular nucleotide selectivity profile. To our surprise, while the *in vitro* activity for the Shortloop mutant enzymes was low, we detected FCD and FUD formation along with FAD by LC-MS in all the cell lysates containing the Shortloop enzyme plasmids (Figure 4C). These results further strengthen the loop’s role in ATP selectivity within the cell.

### The *E. coli Loopswap* strain produces FAD nucleobase analogues

The *ribF* gene encodes the wild-type FAD synthetase enzyme in *E. coli*. We substituted *ribF* with the altered gene sequence that would result in the *LoopSwap* gene product and created the *E. coli* MG1655 K-12 *LoopSwap 155-163 (Loopswap)* strain. LC-MS analysis of the *Loopswap* strain lysate revealed the presence of FAD analogues- FGD, FCD, and FUD (Figure 5A). We then quantified the FAD in the *Loopswap* versus wild-type strain - the wild-type strain contains 1452 ± 411 pM FAD while the *Loopswap* strain contains 918 ± 223 pM FAD, in addition to showing FCD, FGD, and FUD production (Figure 5B, Figure S5A).

**Figure 5.**
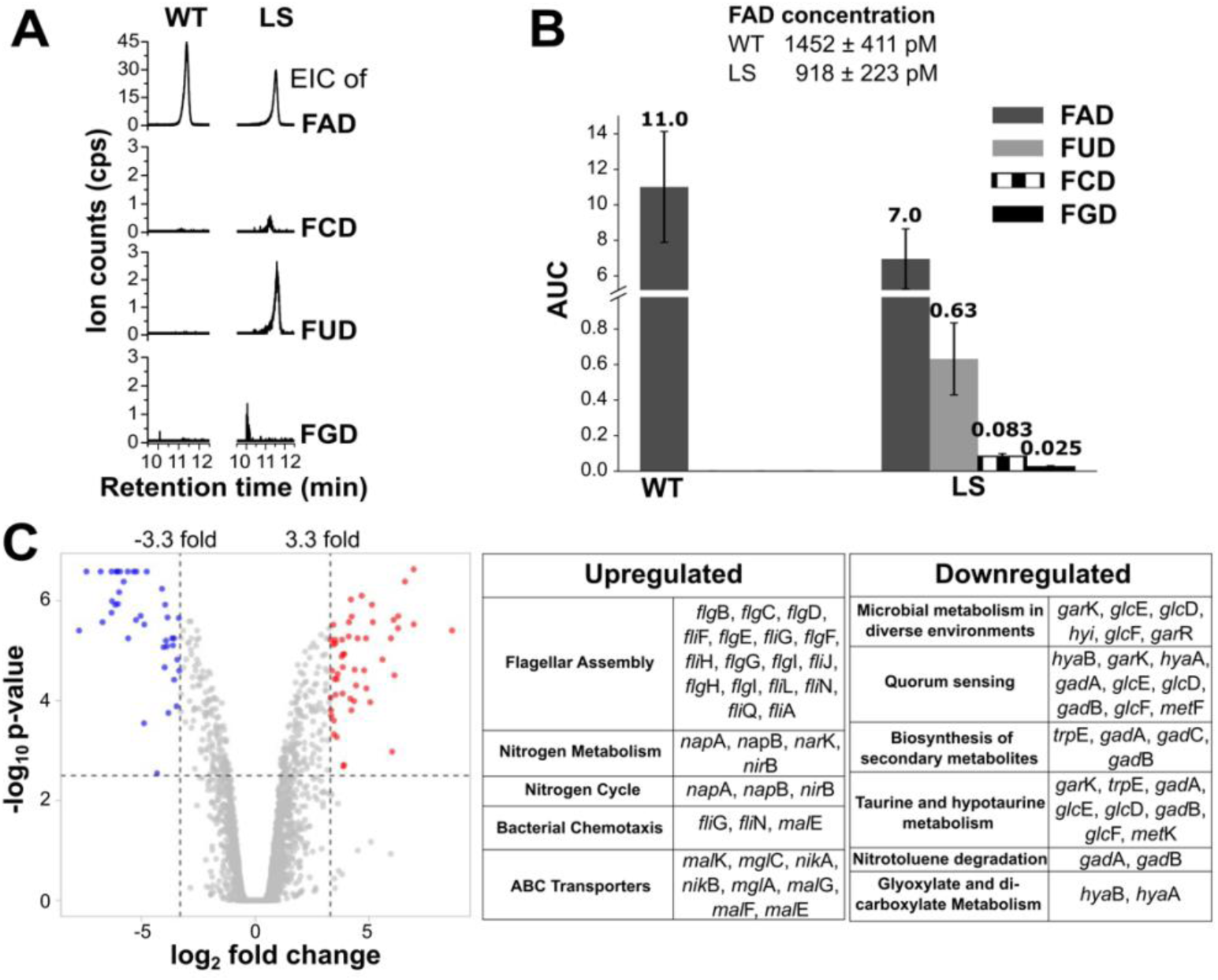
Comparative analysis of the *E. coli* wild-type strain vs *Loopswap* strain. **(A)** EIC of FAD and its analogues in the *E. coli* wild-type and *Loopswap* (LS) strains. **(B)** Area under the curve (AUC) for comparing FAD concentrations in the wild-type and LS strains, and also the FAD analogues. **(C)** Volcano plot generated from RNA sequencing data comparing the wild-type and *Loopswap* strain. A stringent cut-off criterion, log_2_(fold change) above 3.3 and below -3.3, respectively, with p-values below ∼0.003), and their potential functions was set, yielding 92 upregulated (red)/ downregulated (blue) genes.

### Probing the physiology of the *Loopswap* strain

After confirming the presence of the FAD nucleotide analogues in the *Loopswap* strain, we turned our attention to their effects on cellular function. In rich and minimal medium, the *Loopswap* strain displays similar growth to the wild-type strain (Figure 6A-D, S5B). An RNA sequencing (RNA-Seq) analysis was conducted to compare the wild-type and *Loopswap* strains grown in rich medium. With a stringent cutoff of log_2_ (fold change) > 3.3 and < -3.3, we observed that a wide range of physiological functions were altered (Figure 5C). There is an upregulation of genes involved in motility (bacterial flagellar assembly, chemotaxis) and nitrogen metabolism, and downregulation of sulfur (taurine and hypotaurine) metabolism, among others (Figure 5C).

**Figure 6.**
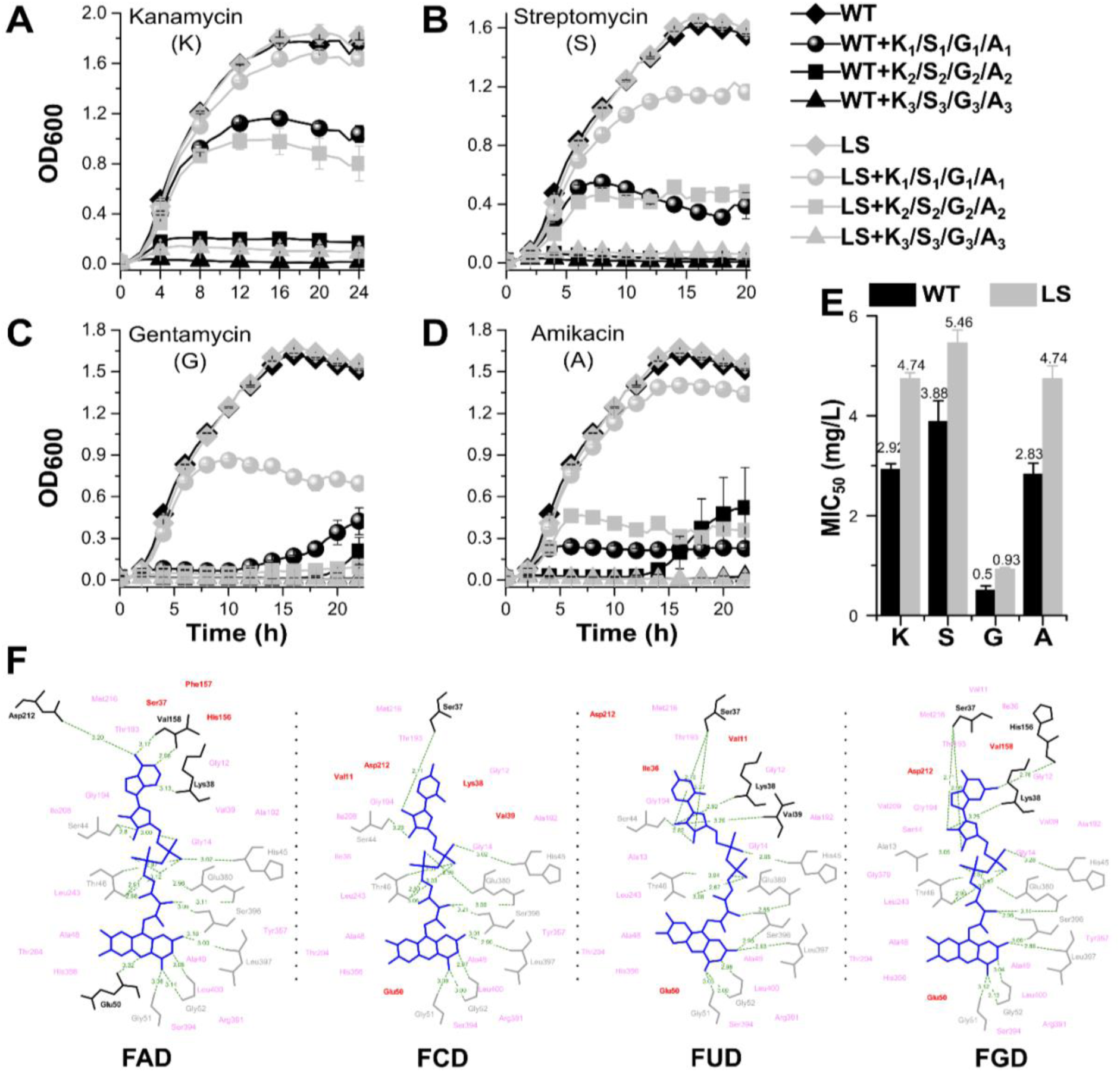
The *Loopswap* strain exhibits antibiotic tolerance against aminoglycosides kanamycin, streptomycin, gentamicin, and amikacin. **(A, B, C, D)** Growth curve of *E. coli* wild-type and *Loopswap* (LS) strains in presence of Kanamycin (K)*, Streptomycin (S)**, Gentamycin (G)^&^, Amikacin (A)^#^. K*-Kanamycin (K1= 2.5, K2= 5, K3= 10 µg/mL), S**-Streptomycin (S1= 5, S2= 10, S3= 15 µg/mL), G^&^-Gentamycin (G1= 1, G2= 2.5, G3= 5 µg/mL), A^#^-Amikacin (A1= 2.5, A2= 5, A3= 15 µg/mL). **(E)** MIC_50_ of aminoglycosides for wild-type and LS strains. **(F)** Interactions of FAD and its analogues docked with fumarate reductase flavoprotein subunit show a favourable fit (UniProt ID: P00363). Differing hydrogen bonding interactions are highlighted in black while differing hydrophobic interactions between analogues are highlighted in red. Common hydrogen bonds are shown in grey, while common hydrophobic interactions between analogues is shown in pink.

### The *Loopswap* strain exhibits decreased antibiotic sensitivity

As nitrogen and sulfur metabolism are closely associated with cellular redox regulation, we investigated whether oxidative stress affects the *Loopswap* strain and how it differs from the wild-type strain. The wild-type and the *Loopswap* strains were grown in the presence of H_2_O_2_ to induce oxidative stress - their growth was suppressed to a similar extent with 2 mM H_2_O_2_, and 5 mM H_2_O_2_ was the lethal concentration (Figure S6A). Next, we studied their growth with a representative set of antibiotics as the source of oxidative stress.^41^ Interestingly, we noticed that *Loopswap* strain grew better than the wild-type when treated with the aminoglycoside class of antibiotics (kanamycin, streptomycin, gentamycin, and amikacin), whereas colistin, ciprofloxacin, trimethoprim, rifampicin, piperacillin, ampicillin, and tetracycline affected both strains equally (Figure 6A-D, S6B-H). To confirm that the aminoglycoside tolerance is indeed caused by the swapping of the loop, we subjected the *Loopswap*-Kan strain (which was an intermediate strain with kanamycin resistance cassette while creating the *Loopswap* strain) to all the oxidative stress agents and obtained identical results (Figure S7). Following this, an extensive antibiotic MIC survey was done for the *Loopswap* and wild-type strains to assess their sensitivity profile. The *Loopswap* strain has a ∼1.6x MIC_50_ value as compared to the wild-type strain for the aminoglycosides, otherwise, both wild-type and *Loopswap* strains were found to have similar MIC_50_ values for all other tested antibiotics (Figure 6E, S6I).

### Metabolomic and computational studies to probe how FAD analogues perturb metabolism and their potential interactions with *E. coli* proteins

The *Loopswap* strain produces FAD and its analogues, so the decreased aminoglycoside sensitivity could be a result of unprecedented enzyme-FAD analogues interactions within the cell leading to altered metabolism. Untargeted metabolomics showed that riboflavin metabolism is downregulated and lower levels of FAD in the *Loopswap* strain as compared to the wild-type strain. Amino acid, nitrogen, sugar, and nucleotide metabolism appear to be perturbed in the *Loopswap* strain (Table S9, Figure S5 C, D, E).^42–44^ The common metabolite that is upregulated across 10 out of 25 altered pathways is glutamate, which has been implicated in oxidative and antibiotic stress tolerance (Table S9).^45–47^

We also launched an extensive computational analysis to examine the antibiotic sensitivity phenotype from the point of view of unprecedented enzyme-FAD analogues interactions. First, we prepared the “*E.coli* K-12 MG1655 protein structural master database”, which is an exhaustive compilation of all *E.coli* protein structures as described in the methods section (Figure S9, Table S3). This database was used to dock FAD and its nucleotide analogues to all *E. coli* proteins and compare their binding affinities. First, we conducted docking studies of the FAD analogues with the FAD-utilizing enzyme glutathione reductase whose activity has been experimentally validated with the FAD analogues (Figure S10).^48,49^ We found that FUD and FGD have similar binding energy as FAD. Additionally, FGD, FCD, and FUD have very similar docking poses as FAD within the glutathione reductase active site. Thus, it is plausible that the FAD analogues may act as potential cofactors for other FAD-binding enzymes. We commenced our proteome-wide docking analysis of FAD and its analogues by validating our docking approach with FAD-binding proteins (as described in the methods section) and found it to be satisfactory (Figure S11A-C, Table S4).

Next, we docked the 74 FAD-binding proteins in the master database with FAD, FCD, FGD, and FUD, and calculated the value ΔA (FAD analogue) = A (FAD analogue) – A(FAD), where A is the protein-ligand binding energy. Docking results from the 11 FAD binding proteins with holo structures resulted in ΔA values between -1.04 to 2.61 kcal/ mol, indicating similar binding (Table S5). We found that the binding poses of FAD and the FAD analogues are similar, with comparable interactions around the catalytic isoalloxazine ring, while the difference in affinity arises from the interactions with the different nucleobases attached to the ligands shown using LigPlot+^50^ (illustrative data with fumarate reductase flavoprotein shown in Figure 6F). We extended this computation of binding affinities to all *E. coli* FAD-utilizing proteins (apo structures from the PDB and modelled structures from AlphaFold DB, full list in Table S6), and the results indicate that 78% FAD-binding proteins bind similarly to the FAD analogues, with ΔA values ranging from -2 to 2 kcal/mol (Figure S12). Our docking studies show that FCD, FGD and FUD bind better to some of the FAD-binding proteins as compared to FAD, and this may lead to some functions being performed differently in the *Loopswap* strain as compared to the wild-type strain (Figure S12).

Finally, to test the possibility that the FAD analogues bind to non-FAD binding proteins causing unprecedented downstream effects, we docked FAD and the FAD analogues with the master database (as described in the methods section). Interestingly, we found 7 candidate proteins that bind stronger to the FAD analogues as compared to FAD, which may be interesting to test experimentally (Figures S13, S14, Tables S7, S8). Specific examples include the outer membrane porin F (OmpF) and the Tryptophan-specific transport protein which are also among the highest upregulated/ downregulated genes in the RNA-Seq analysis (Figure S14). The significance and metabolic role of the protein hits in decreasing aminoglycoside sensitivity is unexplored and requires further molecular-level investigations.

## DISCUSSION

In this study, we explored and altered the nucleotide specificity of the *E. coli* FAD synthetase enzyme to create a range of mutants that can produce FAD and its nucleotide analogues *in vitro* as well as within cells. Surprisingly, we find that the intracellular production of FCD, FGD, and FUD is linked with decreased sensitivity towards aminoglycoside antibiotics.

Firstly, *in vitro* reactions with the wild-type bifunctional *Ec*FADS revealed that its two functional domains, riboflavin kinase and FMNAT, display a difference in the choice of the nucleotide co-substrate. Our studies show that the *E. coli* riboflavin kinase domain converting riboflavin to FMN is specific for ATP and dATP, whereas the FMNAT domain converting FMN to FAD can accept ATP, CTP, UTP, dATP, and dCTP, and can synthesize the corresponding FAD analogues (Figure 2C-F). This is intriguing as the conversion of riboflavin to FMN involves a phosphate transfer and thus, in principle, the reaction is chemically indifferent to the identity of the nucleotide/ phosphate source, whereas the FMN to FAD conversion introduces the nucleotide, and thus its selectivity is important. However, we observed that in spite of the *in vitro* promiscuity of the FMNAT domain, FAD synthetase is specific for ATP within the cell and produces only FAD under native or heterologous expression conditions (Figure S1F). This may be attributed to the cellular ATP concentration being the highest among all nucleotides or to other cellular factors that regulate the entry/ availability of ATP as the co-substrate.

Predicting the amino acid residues that dictate adenine nucleobase specificity and altering them to make the enzyme promiscuous was challenging as their interactions appear to be via the amide backbone.^6,51^ Based on interaction energy calculations and existing literature, mutations at the G23 and S165 positions in the FMNAT domain were conducted, however, G23K and S165K mutants resulted in specificity for ATP accompanied by an overall decrease in activity, and the G23S mutant produced an inactive protein (Figure S2). This suggested that these residues do alter the flexibility of the active site in a significant manner.

Our structural alignment search provided significant insights - notably, we found that the FMNAT domain aligns structurally with cytidylyltransferases with a loop around the nucleobase region that is different.^40^ Swapping the FMNAT loop with that of the glycerol-3-phosphate-cytidyltransferase increased promiscuity for the various nucleotides, that is, overall, the *Loopswap* mutant enzyme produced FGD, FCD, FUD, and dFAD with higher efficiency than the *E. coli* FAD synthetase (Figure 3B and 4B). Also, the overall efficiency of FAD production was retained *in vitro* while the activity with GTP was greatly enhanced as compared to the wild-type enzyme (Figure 4B). Our hypothesis is as follows - as the cellular concentrations of nucleotides in *E. coli* follow the order ATP > GTP > UTP >CTP, the original loop sequence may have enforced ATP binding by a concentration-dependent tuning of its binding affinity.^37^ The swapping of the loop may now allow for lower concentrations of nucleotides to bind and act as substrates for the FMNAT reaction.

Probing the length of the loop sequence by creating the Shortloop enzyme mutants 2-7 additionally confirmed that the loop is an important element in nucleotide choice (Figure 4B, 4C, S4). Another interesting result was that the heterologous expression of the *Loopswap* and *Shortloop* mutants in *E. coli* led to the formation of FCD, FUD, and FGD along with FAD within the cell, as compared to the strict specificity of FAD production with the wild-type FAD synthetase enzyme or an empty vector (Figure 3C, 4C). These observations collectively substantiate the role of the FMNAT loop sequence in *E. coli* FAD synthetase, in its complete structure and sequence, for ATP specificity within the cell. The QTFCEGGV sequence of the FMNAT loop appears to be variable whereas the RIS sequence shows high conservation across organisms, as illustrated by a position-specific substitution matrix (Figure S8). We also find that >60% representative proteins binding ATP, ADP, AMP, CoA, FAD, NAD, SAM and their analogues show the presence of such loops (Table S2).^51^

Finally, to understand the physiological effect of the loop, we generated the *E. coli Loopswap* strain where the wild-type FAD synthetase gene is substituted with the *Loopswap* - FAD synthetase gene. The *Loopswap* strain (i) produces ∼63% FAD as compared to the wildtyp strain, and (ii) produces the FAD analogues (Figure 5A, B). Interestingly, it exhibited a phenotype of 1.6x aminoglycoside antibiotic resistance as compared to the wild-type strain (Figure 6E). This could occur due to the lower FAD content in the *E. coli Loopswap* strain which results in lowering the proton-motive force, a metabolic state that is found in persisters that demonstrate higher antibiotic tolerance.^52^ However, the *E. coli Loopswap* strain appears to have a similar overall growth as compared to the wild-type strain in rich and minimal media, hence this is an unlikely possibility (Figure S5B, S6 A-H, S7A-C, G-I). Thus, we hypothesize that the FAD analogues may play a role similar to FAD in the cell, or be potentially involved in other metabolic pathways. Metabolomic and RNA-Seq studies show large-scale perturbations in primary metabolism between the *Loopswap* and wild-type strains, and strongly implicate a connection of the decreased antibiotic sensitivity with cellular glutamate levels.^45,47^ Computational studies suggest that the FAD analogues can bind to a wide range of enzymes in the cell, which could cause a perturbation of overall metabolism leading to decreased sensitivity to aminoglycoside antibiotics.

## CONCLUSION

In conclusion, this study presents a structural and functional analysis of *Ec*FADS, with a focus on elucidating the determinants of nucleotide selectivity. Engineering a cytidyltransferase-derived loop into the nucleotide-binding pocket enabled broadening of the enzyme’s substrate specificity to accommodate alternative nucleotides. The resulting *E. coli Loopswap* strain synthesizes FAD and its noncanonical nucleotide analogues, while exhibiting increased tolerance to aminoglycoside antibiotics. These findings suggest that targeted modification of the loop region proximal to the nucleobase-binding site can modulate nucleotide specificity in adenylating enzymes, and can be applied to the synthesis of other noncanonical redox cofactors (REF). This study advances our understanding of FAD biosynthesis and its metabolic roles, including its unexpected link to antibiotic sensitivity. Finally, it highlights the potential for developing bio-orthogonal redox systems in synthetic biology via cellularly produced FAD nucleobase analogues, thus expanding the biochemical toolkit for metabolic engineering and biotechnological innovation.

## METHODS

### Cloning and purification of the *Ec*FADS enzyme and mutants

The cloning, protein expression, and purification of the *Ec*FADS enzyme and its mutants was conducted using standard molecular biology and protein purification techniques noted in the supplementary materials methods section.

### Activity assay of *Ec*FADS and mutants

The activity of *Ec*FADS was optimized for the presence and absence of dithiothreitol (DTT), and varying concentrations of MgCl_2_, and time. The activity assay conditions optimized for *Ec*FADS enzyme and the mutants with NTPs and dNTPs were: Tris-HCl (100 mM, pH 8.0), riboflavin or FMN (200 μM), ATP (3.5 mM), DTT (20 mM), MgCl_2_ (3 mM) and 5 μM of the enzyme. The same assay conditions were used with physiological amounts of other NTPs and dNTPs for *Ec*FADS and mutants as described in the supplementary materials methods section.

### Homologous recombination to generate the *Loopswap* strain

*E. coli* K-12 MG1655 wild-type-Kan and *Loopswap* -Kan mutant strains were generated using recombination by the λ Red recombineering system.^53^ The kanamycin resistance cassette was excised using FLP recombinase to generate the *E. coli* K-12 MG1655 wild-type and *Loopswap* strains, leaving a scar sequence.^54^ Details of the protocol for homologous recombination and kanamycin cassette removal are described in the supplementary materials methods section.

### LC-MS analysis for detection of FAD analogues in cell lysates

*E. coli* BL21(DE3) strain transformed with empty pET28a(+) plasmid vector, *Ec*FADS-pET28a (pLP001), or *mutant*-pET28a constructs were grown, lysed, and the supernatant was subjected to LC-MS analysis as described in the supplementary materials methods section.

Wild-type and *Loopswap* strain lysates were analysed on LC-MS as described in the supplementary materials methods section.

### RNA sequencing and data analysis

Sample preparation for RNA sequencing was conducted using previously published methods, mentioned in detail in the supplementary methods section.^55^ RNA extraction, sequencing, and preliminary data analyses as per the reference transcriptome of *Escherichia coli* were performed by Strand Life Sciences (India). RNA sequencing data was analysed on DAVID^56^ (Database for Annotation, Visualization and Integrated Discovery, v6.8) for functional enrichment and pathway analysis using the Kyoto Encyclopedia of Genes and Genomes pathway.

### Growth assay of the wild-type and *Loopswap* mutant strain under antibiotic stress

The primary culture for each strain was grown overnight from glycerol stock in LB media at 37 °C, 180 rpm. Then, in LB media, secondary culture was grown under the stress of various antibiotic classes (Aminoglycosides, Beta-lactams, Fluoroquinolones, Antifolates, Polymyxins, Ansamycins, and Tetracyclines) with varying concentrations and the growth was monitored over 24-48 h. The MIC_50_ determination was conducted using previously published methods.^55^ Details of all these procedures are noted in supplementary materials methods section.

### Untargeted Metabolomics using mass spectrometry

Wild-type and *Loopswap* mutant MG1655 strain lysates were analysed on LC-MS using the Information Dependent Acquisition (IDA) mode (parameters detailed in supplementary section). The XCMS^57^ Online tool was used for pairwise analysis of metabolites in wild-type and *Loopswap* mutant strain, followed by a pathway enrichment analysis using LCMS Functional analysis tool in MetaboAnalyst 6.0.^58^ Details for this procedure are in the supplementary materials methods section.

## Computational methods

### Finding structural analogues of *Ca*FADS

We used the Distance-matrix ALIgnment (DALI)^39^ server to search the structural homologs of *Ca*FADS, the detailed parameters of which are mentioned in the supplementary methods section.

### Bioinformatic analysis for generation of *Shortloop* mutants

A position-specific substitution matrix (PSSM) was generated using Biopython for the loop sequence of *Ec*FADS protein with a set of 5000 homologous proteins. *Shortloop* mutants designed based on the PSSM were modelled with Phyre2.^35^

### Compiling of the *E. coli* K-12 MG1655 protein structural database

We constructed a structural database (crystal/ cryo-EM/ AlphafoldDB) of all *E. coli* proteins using a protocol we designed, detailed steps of which are described in the supplementary methods section. The compiled *E. coli* K-12 MG1655 protein structural master database contained 4382 proteins (out of 4403 total known *E. coli* proteins) (Table S3, schematic in Figure S9). Out of these, 1116 had crystal structures. Among these structures, if the PDB dataset (Dataset 2) contained apo and holo versions of the same protein (using a “non-polymer bound ligand query” for each PDB ids), both were considered with an apo/ holo tag. We found 790 apo and 650 holo tagged versions of the RCSB protein structures (Table S3 - Extended Datasets) were added to obtain a total of 4706 structures.

### Selecting FAD-binding proteins from the *E. coli* K-12 MG1655 protein structural database

A total of 74 of *E. coli* K-12 MG1655 FAD-binding apo- or holo- protein structures were obtained from Uniprot through a keyword-based search (Table S6). Search parameters are given in the supplementary methods section.

### Docking protocol and its validation

Blind docking was performed using DiffDock-L^48^ for *E. coli* K-12 MG1655 protein structural master database and validated on FAD-binding and non-FAD-binding *E. coli* proteins (Figure S11A, B, C). Docking and validation method details are given in the supplementary methods section.

### Interaction of known FAD-binding proteins with FCD, FUD, and FGD

The docking scores of FAD, FCD, FGD, and FUD from the 74 FAD-binding structures from the *E. coli* K-12 MG1655 protein structural master database were compared and the difference in affinity (ΔA) was computed between FAD and each of its analogues (Figure S12).

### Interaction of all proteins in the *E. coli* K-12 MG1655 protein structural master database with FAD, FCD, FUD, and FGD

The docking scores of FAD, FCD, FGD, and FUD with non-FAD-binding structures from the *E.coli* K-12 MG1655 protein structural master database were computed. Structures within the *E. coli* K-12 MG1655 protein structural master database with less than 30 amino acids and more than 1022 amino acids were not considered for docking based on DiffDock’s criteria. Hence, this exercise was conducted for 4555 structures. Each ligand-protein pair yielded 10 docking scores. The analysis involved the following. (1) Ranking ligand poses by binding energy and confidence score. (2) Filtering proteins that have a confidence score > -1.5 and binding affinity < -11 kcal/mol. (3) Filtering proteins with fewer than five FAD poses among the top ten ranked ligand poses and confidence scores. (4) Identifying proteins common to these filtered lists (Table S7). (5) Evaluating whether these proteins are associated with aminoglycoside resistance based on existing literature. The docked structures with FAD, FCD, FGD, and FUD from the proteins listed in Table S7 are shown in Figure S13.

## ACKNOWLEDGEMENTS

Financial support was provided by the Ministry of Science and Technology, Govt. of India – SERB Core Research Grant CRG/2019/003270 for this research to A.B.H. and for salary to A.S. We acknowledge UGC for providing fellowship to S.M. The authors acknowledge the DST fund for Improvement of S&T Infrastructure (SR/FST/LSII-043/2016) to the IISER Pune Biology Dept for the Biological Mass Spectrometry Facility and the IISER Pune NMR and MS facilities and their managers. The support and the resources provided by ‘PARAM Brahma Facility’ under the National Supercomputing Mission, Government of India at the Indian Institute of Science Education and Research (IISER) Pune are gratefully acknowledged. We thank DBT, Govt. of India (Grant no. BT/PR34215/AI/133/22/2019) to A. M. for partial funding of the computational resources used in this work. The authors are grateful to Hemalatha Balaram for critical discussions, and Siddhesh Kamat and Aakash C. for their help with mass spectrometric metabolomic analysis.

## Notes

### Competing Interest Statement

The authors have declared no competing interest.

### Summary of Updates

An additional metabolomics analysis has been conducted, reported in the Results section, and analyzed in the Discussion section. Supplemental files have been updated.

