## Supplemental methods and figures for "The biosynthesis of FAD nucleotide analogues in *Escherichia coli* alters its sensitivity to aminoglycoside antibiotics"

### MATERIALS AND METHODS

#### Experimental Methods:

##### Materials

Nucleotide triphosphates (NTP) and 2'-deoxy nucleotide triphosphates (dNTPs) were obtained from Jena Biosciences and Sigma. Reagents and chemicals for the enzymatic assays were purchased from RANKEM Chemicals, TCI Chemicals, and Merck. All reagents and components of media used in bacterial cultures were ordered from TCI Chemicals, HiMedia, and SRL Chemicals. DSS Takara Bio USA supplied Taq polymerase, Dpn1, and the PrimeSTAR GXL DNA polymerase. Purification kits for DNA and plasmid were obtained from Agilent and Qiagen. All solvents for high-performance liquid chromatography (HPLC) and liquid-chromatography mass spectrometry (LC-MS) analysis were obtained from Merck, SD Fine Chemicals, and RANKEM Chemicals. All primers have been purchased from Eurofins as consolidated in Table S10.

##### Cloning and purification of *Escherichia coli* flavin adenine dinucleotide (FAD) synthetase (*Ec*FADS) and mutants of the flavin mononucleotide adenylyltransferase (FMNAT) module

The cloning of the *ribF* gene for *Ec*FADS was done in a pET28a(+) plasmid between BamHI and NdeI restriction sites using restriction-free cloning.[1] The first round of PCR was done for amplification of the *ribF* gene from *E. coli* genomic DNA using the first set of primers (*Ec*FADS_F1 and *Ec*FADS_R1; Table S10). The second round of PCR added flanking sequences at terminals of the gene that are complementary to the insertion sites of the pET28a vector. This step was done using the second set of primers (*Ec*FADS_F2 and *Ec*FADS_R2; Table S10). The gene obtained in the second round of PCR, megaprimer, was used to amplify the whole plasmid vector with the gene inserted between the restriction sites mentioned above. The parent plasmid was digested using Dpn1, and chemically competent *E. coli* DH5α cells were transformed with the amplified nicked plasmid, followed by the isolation of the cloned vector. For the cloning of mutants, the wild-type *ribF*-plasmid vector was employed as a template, and the list of the corresponding primers is given in Table S10. These primers contain the altered codon at the site of mutation. The mutants were cloned by PCR in two rounds. The first round of PCR was done with the corresponding forward primer of the mutant and the T7 terminator universal primer (Table S10). It amplified the fragment of plasmids from the T7 terminator site to the mutation site. This fragment, called megaprimer, was utilized in the other round of PCR to amplify the whole plasmid introduced with mutation, which was transformed into DH5α chemically competent cells followed by plasmid isolation.

The *ribF* or the mutant gene containing isolated plasmids were introduced to the chemically competent *E. coli* BL21(DE3) cells. Overnight grown primary culture was utilized to grow the secondary culture in autoclaved Luria Bertani (LB) broth (1 L) with kanamycin (35 μg/mL), followed by the growth at 37 °C and 180 rpm. Protein overexpression was induced at 28 °C by adding 0.1 M isopropyl-β-D-1-thiogalactopyranoside (IPTG) once the culture's $OD_{600}$ reached to a range of 0.6 to 0.8. The culture was then grown for 8 hours, followed by pelleting through centrifugation (6000 rpm for 15 min) and subsequently pellet was stored at -80 °C. The Ni-NTA affinity column (GE Healthcare) chromatography was used to purify the N-terminally hexa-histidine-tagged protein. This method utilized the Äkta pure FPLC system from Cytiva Lifesciences (formerly GE Healthcare). The pellet was lysed in the 20 mL lysis buffer (50 mM Tris-HCl pH 8.0, 0.025% β-mercaptoethanol, 20 mM imidazole, and 300 mM NaCl) on sonicator (1 sec pulse-on 3 sec pulse-off for 20 min at 60% amplitude) followed by centrifugation (18000 rpm for 25 min). Initially, Ni-NTA column was equilibrated using lysis buffer followed by supernatant loading. The column was washed with lysis buffer followed by wash buffer (50 mM Tris-HCl pH 8.0, 0.025% β-mercaptoethanol, 100 mM imidazole, and 300 mM NaCl). Elution of the protein was accomplished using the elution buffer (50 mM Tris-HCl pH 8.0, 0.025% β-mercaptoethanol, 500 mM imidazole, and 300 mM NaCl). To determine protein concentration, the Bradford assay was performed with standard Bovine serum albumin (BSA). An identical procedure was used for all *Ec*FADS variants. The protein's molecular weight and purity were determined by SDS-PAGE gel electrophoresis.

**Activity assay of *Ec*FADS and its mutants**

The activity of *Ec*FADS was optimized using Tris-HCl (100 mM, pH 8.0) buffer, riboflavin (200 μM), ATP (3.5 mM), and the enzyme (5 μM) with or without 20 mM DTT (dithiothreitol) and varying concentrations of $MgCl_2$ (0-10 mM). For different divalent metal ions, 3 mM concentration of $M^{2+}$ ion was used instead of $MgCl_2$ in the presence of 20 mM DTT while keeping all other components the same. The time optimization was done with 3 mM $MgCl_2$ for several time points in the presence of 20 mM DTT.

The final assay for the activity of *Ec*FADS and the mutants with NTPs and dNTPs was as follows: Tris-HCl (100 mM, pH 8.0), riboflavin or FMN (200 µM), ATP (3.5 mM), DTT (20 mM), $MgCl_2$ (3 mM) and 5 µM of the enzyme in final concentration. The activity with other NTPs (UTP, CTP, and GTP) and all dNTPs (dATP, dTTP, dCTP, and dGTP) was checked similarly at 3.5 mM concentration. All the reactions were performed for 6 h at 37 °C. Thin-layer chromatography (TLC) was used to visualize the fluorescent flavins excited at 365 nm. The mixture of water, acetic acid, and n-butanol was used in the ratio 5:3:12 as a solvent system.

For *Ec*FADS activity, the cellular concentrations of NTPs - 3.5 mM ATP, 1.5 mM GTP, 0.8 mM UTP, 0.5 mM CTP, 0.2 mM dATP, or 0.2 mM dCTP - were used, keeping the other components the same.[2] Aliquots at various time points were taken and observed on TLC and by HPLC. An Agilent 1260 Infinity-II series HPLC linked with a UV-Vis diode array detector was utilized to assess enzymatic reactions. The analysis used a Phenomenex Gemini reversed-phase column of 250 x 4.6 mm dimensions with a 5 µm particle size of NX-C18. The solvent A was ammonium acetate buffer (10 mM, pH 6.0), and solvent-B was methanol. A method spanning 60 minutes at 0.5% flow rate was applied, 0-15 min 5% B, followed by a gradient increase to 80% B from 15-40 min, followed by a gradient decrease to 5% B from 40-55 min, and finally, 55-60 min at 5% B. A shorter method spanning 30 minutes with the same solvent system and flow rate was also used in some runs, 0-2.5 min at 5% B, followed by a gradient increase to 80% B from 2.5-20 min, followed by a gradient decrease to 5% B from 20-27 minutes, and finally holding at 5% solvent B from 27-30 min. The percent of FMN consumed in each reaction was measured by area under the curve (AUC) of FMN peak in the sample relative to the AUC of FMN in the control sample.

**Heterologous expression of *Ec*FADS or mutants for lysate analysis of FAD and its analogues by LC-MS**

50 mL LB broth was inoculated with the 1% overnight grown *E. coli* BL21(DE3) strain transformed with empty pET28a(+) plasmid vector, *Ec*FADS-pET28a (pLP001), or *mutant*-pET28a (pAS025, pAS026, pAS027, pAS028, pAS029, pAS030 and pAS031) constructs and the culture was grown with kanamycin (35 µg/mL) at 37 °C followed by continuous shaking on 180 rpm. Only the *E. coli* BL21(DE3) culture was grown without kanamycin. We did not induce the culture with IPTG and instead relied on the leaky expression of the pET28a plasmid to minimally perturb the cellular metabolism. After 14-16 h, the culture was centrifuged at

5500 rpm for 15 min at 4 °C resulting in a pellet. 2 mL of 90 % methanol was used for the resuspension of the pellet followed by sonication at 100 % amplitude for 3 min with 1s pulse on and 1s pulse off. The final supernatant was obtained by centrifuging the lysate at 4 °C with 5500 rpm for 15 min, followed by the filtration via 0.22-micron filters. The supernatant was used directly or concentrated on high vacuum for analysis on LC-MS.

A Mass spectrometer (SCIEX X500R-QTOF) was employed for LC-MS analyses, where a UHPLC (Exion-LC series) was used to separate the flavins. The flavins were detected in negative mode at a temperature of 400 °C. The parameters are as follows: ion spray voltage was -4500 V, the de-clustering potential was -80 V and collision energy was 5V. The SCIEX-OS Explorer software facilitated data acquisition, processing, and the generation of metrics such as extracted ion chromatogram (EIC), total ion chromatogram (TIC), and the m/z values of targeted molecules. The analysis used a Phenomenex Gemini reversed-phase column of 250 x 4.6 mm dimensions with a 5 µm particle size of NX-C18. The solvent-A was ammonium acetate buffer (10 mM, pH 6.0), and methanol was used as solvent-B. A method spanning 32 minutes was used at the flow rate of 0.5 mL/min, starting with 20% of solvent B at 0 minutes, maintaining this ratio until 1 minute, a gradient increase to 60% of solvent B until 20 minutes, followed by a gradient increase to 100% of solvent B until 23 minutes, then a return to 20% solvent B until 24 minutes, and finally, maintaining at 20% solvent B until 32 minutes.

**Homologous recombination of wild-type-Kan and *Loopswap*-Kan mutant**

Both *E. coli* K-12 MG1655 *wild-type*-Kan and *Loopswap*-Kan (LS-Kan) mutant strains were generated using recombination by the λ Red recombineering system.[3] *Ec*FADS or *Loopswap* mutant expressing genes were amplified from cloned pET28a(+) plasmid using following primers (forward primer1 5'-GACCGCTGTACAAGGTATACTCGGACGATTTTCACTGTTTTGAGCCAGACATGAAGCTGATACGCGGCATACATA-3' and reverse primer1 5'-ACTTCGAAGCAGCTCCAGCCTACACTTAAGCCGGTTTTGTTAGCCCAAAA-3'), where forward primer has 50 extra complementary bases for *E. coli* genome upstream to *Ec*FADS expressing gene (*ribF*) and the reverse primer has 25 extra complimentary bases for pKD4 vector. Further, the amplified gene was directly stitched to *kan* cassette (including FRT sites) from pKD4 vector by taking the amplified *Ec*FADS and *Loopswap* gene along with forward primer1 as forward primer and the reverse primer2 (5'-TCACTCATCAGATTCTCGGTTCCGTATTTCGGTTTGATTACATAACAGGCATATGA

ATATCCTCCTTAGTTCCTA-3') which has 50 complementary bases to the *E. coli* genome downstream to *ribF* gene. The 50 complementary bases to the *E. coli* genome upstream and downstream to the *ribF* gene were used for homologous recombination.

For the homologous recombination, 5 mL of *E. coli* K-12 MG1655 transformed with pKD46 (which expresses λ red recombinase and is temperature sensitive) was grown overnight at 28 °C with ampicillin (100 μg/mL) in Luria Bertani (LB) media. 30 mL of secondary culture was grown with 1% overnight grown primary culture, 100 μg/mL Ampicillin, and 0.2 % arabinose in LB media for 3 h at 28 °C (till $OD_{600}$ reached 0.3-0.5). The cells were pelleted down by centrifugation (4000 rpm for 10 minutes), followed by resuspension in 10 mL of autoclaved milli-Q water already kept on ice. Then, the cells were again centrifuged, followed by resuspension in 600 μL of autoclaved chilled milli-Q water. Then, the resuspended cells were transformed with the PCR stitched DNA fragments (*Ec*FADS-Kan or *Loopswap*-Kan) by electroporation. For recovery, the electroporated cells were grown at 37 °C for 5 h in 1 mL LB media, followed by spreading on an LB agar plate with 35 μg/mL kanamycin antibiotic and incubation at 37 °C. The obtained colonies were screened by colony PCR using the following primers (forward primer2 5'-CTCAATCGCCGGTTAACCTT-3' and reverse primer3 5'-CGGCAAATTCAGGGTTGATTTATAG-3'), and the amplified gene fragment including *kan* cassette was confirmed by sequencing.

The *kan* cassette was removed by transforming the mutant strain with pCP20 plasmid, which has yeast Flp recombinase which detect and cut at the FRT sites of DNA.[3,4] Both the strains were grown in 5 mL LB media till $OD_{600}$ reached 0.4 - 0.5. The cultures were centrifuged (4000 rpm for 10 minutes) at 4 °C, followed by discarding the supernatant. The pellet was again resuspended in 0.1 M calcium chloride (3 mL) and kept at 4 °C for 10 minutes. The culture was again centrifuged at 4 °C, and the supernatant was discarded, followed by resuspending it in 0.1 M calcium chloride (1 mL). This time, the resuspended cells were kept at 4 °C in calcium chloride for 30 minutes to make the cells chemically competent. These competent cells were transformed with PCP20 plasmid by heat sock at 42 °C for 1 minute, and then the addition of 0.3 mL LB media was done. The transformed cells were kept shaking at 30 °C for 4 hours, and the grown cells were spread on LB agar with ampicillin antibiotic (100 μg/mL) and kept at 30 °C till colonies were observed on the plate. To remove the temperature-sensitive pCP20 vector from the strains, the obtained colony was grown in LB media at 42 °C for 5-6 hours. The culture was diluted $10^{-6}$ times, and around 100 μL was spread on LB agar, incubating overnight at 37 °C. Four colonies were picked up and streaked on four different LB agar plates as follows: (i) only LB agar plate at 37 °C, (ii) LB agar plate with ampicillin (100 μg/mL) at 30 °C, (iii) LB agar plate with kanamycin (35 μg/mL) at 37 °C, and (iv) LB agar plate with ampicillin (100 g/mL) and kanamycin (35 μg/mL) at 30 °C. Only the LB agar plate without antibiotics had the grown colonies, confirming the removal of the *kan* cassette and pCP20 vector.

**LC-MS analysis for FAD analogues in the lysate of wild-type MG1655 and *Loopswap* mutant strain**

Inoculation of 50 mL LB broth was done with overnight grown primary culture of both the strains, and the culture was grown at 37 °C and 180 rpm. After 14-16 h, the culture was centrifuged at 5500 rpm for 15 minutes at 4 °C resulting in a pellet. 2 mL of 90 % methanol was used for the resuspension of the pellet followed by sonication at 100 % amplitude for 3 minutes with 1s pulse on and 1s pulse off. The final supernatant was obtained by centrifuging the lysate at 4 °C at 5500 rpm for 15 minutes, followed by the filtration via 0.22-micron filters. The supernatant was kept at a high vacuum to concentrate it directly for analysis on LC-MS as described previously. The normalised (by $OD_{600}$ =1 and volume =1 mL) EICs of each FAD analogue from both strains were used for the analysis of area under the curve (AUC) where the extracted AUCs of the individual FAD analogue resulted the relative amount of that FAD analogue in the cell (Figure 5B). The FAD analogues in the wild-type strain were not subject for quantitation because of no detection. Further, the FAD concentration was measured using the AUC from normalised EICs using standard curve of AUCs from standard FAD (Figure S7).

**Growth assay of wild-type and its *Loopswap* mutant strain under antibiotic stress**

The primary culture for each strain was grown overnight from glycerol stock in LB media at 37 °C, 180 rpm. Then, in LB media, 200 μL secondary culture was grown under the stress of various antibiotics with varying antibiotic concentrations at a starting $OD_{600}$ of 0.1 in a 96-well plate equipped with a lid, and the plate was incubated aerobically at 37 °C, with ~300 rpm orbital shaking, for 24 to 48 h. $OD_{600}$ readings were taken after every shaking cycle of 1 hour inside a BMG Labtech CLARIOstar® Plus microplate reader.

The $MIC_{50}$ determination was conducted using previously published methods.[5] The primary culture for each strain was grown overnight from glycerol stock in LB media at 37 °C, 180 rpm. Then, in LB media, 200 μL secondary culture was grown under the stress of 2-fold serially diluted antibiotic concentrations at a starting $OD_{600}$ of ~0.05 in a 96-well plate equipped with a lid, and the plate was incubated aerobically at 37 °C, with ~300 rpm orbital shaking, for 20 h. $OD_{600}$ readings for each concentration of antibiotic were recorded on BMG Labtech CLARIOstar® Plus microplate reader. The average from triplicate OD600 readings was done and normalised by taking maximum OD600 as 1 followed by a plot against the log10 value of corresponding antibiotic concentration using OriginPro 9 software. The curve fitting was done as *Analysis > Fitting > Nonlinear Curve Fit > Open Dialog (a dialog box appears) > Function Selection* > select *Category* as *Growth/Sigmoidal* > select *Function* as *Logistic.* The fitting provides the $MIC_{50}$ value.

**RNA sequencing and data analysis**

Sample preparation for RNA sequencing was conducted using previously published methods.[5] Primary culture was set up picking 3 isolated colonies in 3 mL LB as three biological replicates for 17 h at 37 °C, 180 rpm. 1% of saturated primary culture was used to set up 3 mL LB secondary culture for 3 h at 37 °C, 180 rpm to attain the log phase of growth. The 3 mL secondary cultures were pelleted down at 13000 rpm for 2 min. The cell pellet was redissolved in 1mL RNAlater (Invitrogen) solution and Strand Life Sciences (India) performed the RNA extraction, sequencing, and preliminary data analyses. The cell pellets were subjected to lysis and extraction using the Trizol method. The isolated RNA was quantified using qubit and quality assessed using Tapestation. The Kapa Hyperplus kit was used to perform library prep. The library generated was then amplified with minimal PCR cycles to avoid any bias. The amplified library was subjected to clean up using magnetic beads. The expected library size was checked by running a tapestation. A library was considered a pass if it resulted in a right peak at the expected size and yielded enough for sequencing. The libraries were normalized, pooled and sequenced using NovaSeq X plus. 10B flow cell, 300 cycles kit was used to generate 10 million reads per sample with 150bp read length.

Demultiplexed FASTQ files were imported into Strand NGS v4.0 for alignment and analysis. The reads were aligned to the reference transcriptome of *Escherichia coli*, using Strand NGS v4.0. *Escherichia coli* str. K-12 substr. MG1655, complete genome sequences (NC_000913.3). The volcano plot was generated using web-based tool VolcaNoseR and highly upregulated and downregulated genes were filtered out based on $\log_2$(fold change) above 3.3 and below -3.3, respectively, with p-values below ~0.003.[6]

The filtered lists of upregulated and downregulated genes were then submitted separately with the "Official_Gene_Symbol" identifier to DAVID (Database for Annotation,

Visualization and Integrated Discovery, v6.8) for functional enrichment and pathway analysis.[7] The species was set to "*Escherichia coli* str. K-12 substr. MG1655". Then, on the Annotation Summary results page, we select "KEGG Pathway" under 'Pathways' section to identify biological pathways and metabolic processes most affected in the *Loopswap* mutant compared to the wild-type, to correlate the large-magnitude transcriptional changes with the observed phenotypes.

**Untargeted Metabolomics**

Untargeted metabolomic profiling was performed using an information-dependent acquisition (IDA) method in both positive and negative electrospray ionization modes. Source parameters were set as follows: Ion Source Gas 1, 55 psi; Ion Source Gas 2, 50 psi; Curtain Gas, 30; Collisionally Activated Dissociation (CAD) gas, 7; and source temperature, 500 °C. Spray voltages were +5500V and −4500V for positive and negative modes, respectively. TOF MS scans were acquired from 100–1000Da with an accumulation time of 0.1s. Declustering potential (DP) was ±80V with a DP spread of 20V. Collision energy (CE) was ±10V with a CE spread of 5V. IDA was triggered for small-molecule selection with dynamic background subtraction enabled, an intensity threshold of >100 cps, and a maximum of 20 candidate ions per cycle. TOF–MS/MS spectra were acquired over 50–1000 Da with an accumulation time of 0.05s. MS/MS parameters were set to DP ±55V (DP spread 20V) and CE ±25V (CE spread 10V).

WIFF files obtained from LCMS runs were converted to the .mzML format using ProteoWizard- MSConvert tool.[8] Centroiding was performed using OpenMS-PeakPickerHiRes tool, with a signal to noise ratio of 1.0 and MS levels 1 and 2 chosen for peak picking.[9] Centroided .mzML files of wild-type and *Loopswap E. coli* MG1655 samples were processed in XCMS Online for both ionization modes in a pairwise job.[10] Feature detection used the centWave algorithm (ppm 30, peak width 5–5 s, signal-to-noise 10, m/z diff 0.05, noise filter 500). Retention-time correction employed the peakgroups method with loess alignment (bw 5, mzwid 0.05, minfrac 1, span 0.2); alignment parameters were bw 10, minfrac 0.5, mzwid 0.01, minsamp 2. Statistical testing used an unpaired Welch's t-test with post hoc analysis; features with $p < 0.01$ and fold change $\geq 2$ were considered highly significant. The XCMS analysis identified 1071 (negative mode) and 881 (positive mode) features.

XCMS results were separated into up- and down-regulated groups to generate four CSV files (columns: m/z median, retention time median, p-value) for LCMS Functional analysis [LCMS] tool in MetaboAnalyst 6.0.[11] Pathway enrichment was performed using Mummichog 2.0 algorithm with 10 ppm mass tolerance, *E. coli* K-12 MG1655 [KEGG] library, and a p-value cutoff of 0.05 (top 10% peaks).[12]

#### Computational methods:

##### Obtaining the structural model of *Ec*FADS

For sequence alignment, a web-based tool, T-Coffee, was used.[13] All structure visualization and structure alignment were done with PyMOL.[14] The homology modelling was done using a web-based tool, Phyre2 (confidence score of 100%) and MODELLER.[15,16]

##### Finding *Ca*FADS/ *Ec*FADS structural analogues

We used the DALI server to search the structural homologs of *Ca*FADS.[17] PDB90 search was employed to eliminate redundant PDBs with sequence similarity above 90%, and a list is provided which was shorted by z-score of 10 and above (Table S1). The DALI search result showed the glycerol-3-phosphate-cytidyltransferase to have high structural homology with FMNAT domain of *Ec*FADS, except for the PRTEGIS sequence loop adjoining the cytosine. Hence, the corresponding loop sequence of the *Ec*FADS FMNAT domain (QTFCEGGVRIS) adjoining the adenine nucleobase was selected for creating the Loopswap mutant by swapping the two loops.

##### Validation of the loop sequence and generation of *Shortloop* mutants

To get a sense of how often a loop structure appears in the ATP- binding region of a protein, we looked for the presence of such loops in 80 representative PDB structures from the available published list of 985 crystal structures bounded to ATP, ADP, AMP, CoA, FAD, NAD, SAM and their analogues.[18] We defined the loop as at least four consecutive residues present around the nucleobase, and loops that are positioned away from the nucleobase in the crystal structure were not considered. Our findings revealed that 62% of proteins utilizing ATP and its derivatives featured this distinct loop structure with varying sizes (Table S2).

A BLASTp search was done against the reference *Ec*FADS protein sequence, excluding "Gammaproteobacteria" and by setting the 'maximum target sequences' to 5000.[19] The FASTA file from the BLAST output was downloaded, to which the reference protein sequence was added. Multiple sequence alignment was performed using muscle version 3.8.31.[20] Finally, the position-specific substitution matrix (PSSM) was generated using Biopython [21].

**Creating *Shortloop* mutants**

To further create the *Shortloop* mutants, we compared the sequence of the two loops. The IS sequence is strictly conserved, hence the variants QTFCEGIS, QTFCIS, QTFIS, and QTIS were designed, making the loop shorter (PSSM shown in Figure S6). These mutants were modelled with Phyre2[15] in NORMAL mode with 100% confidence in the models. Based on both the loop sequences, since the GIS appeared to be common and structural alignment of these *Shortloop* mutants with glycerol-3-phosphate-cytidyltransferase, QTFCGIS and QTCGIS mutants were also created.

**Molecular Dynamics simulations of the *Ec*FADS mutants**

We conducted docking and MD simulations for *Ec*FADS and its predicted mutants (S165K, G23K, and QTFCGIS). We constructed models for both wild-type *Ec*FADS and its mutants using MODELLER.[16] The AutoDock4 software was used for docking, followed by the molecular dynamics (MD) simulation of the docked complex.[22,23] For the simulation, we used AMBER99SB force field for the protein, and generated general amber force field (GAFF) for ATP, CTP, GTP, and FMN using the antechamber module of AmberTool.[24–26] We began with the quantum mechanical optimizations of ATP, GTP, and CTP using the HF/6-31G* basis set in GAUSSIAN03 software.[27] Subsequently, AmberTool was utilized to compute restricted electrostatic potential (RESP) charges for the atoms within ATP, GTP, CTP, and FMN.[28] The resulting topology and coordinates, produced by AmberTool, was converted into GROMACS format using the amb2gmx.pl program. Molecular dynamics simulations were carried out using GROMACS package (version 4.5).[29] We placed the NTP-FMN bound protein within a cubic box measuring 70 Å in length, and solvated it with TIP3P water molecules.[30] We introduced a solution of 150 mM $MgCl_2$ ions and optimized the system using the steepest descent method.[31] The subsequent steps involved heating the system to 300 K using the Berendsen thermostat with a coupling constant of 0.2 ps under NVT conditions.[32] After this, a 10 ns NPT simulation was performed at 300 K and 1 bar pressure, adjusting the box size by employing the Berendsen thermostat and barostat with a coupling constant of 0.4 ps.[32] Finally, a production run lasting for 100 ns at 300 K was conducted using the Nose-Hoover thermostat.[33] All subsequent analyses were carried out on the trajectory of this production run.

**Compiling of the *E. coli* K-12 MG1655 protein structural database**

We constructed a structural database (crystal/ cryo-EM/ AlphafoldDB) of all *E. coli* proteins using the following protocol.[34,35]

1. The accession IDs of all *E. coli* K-12 MG1655 proteins (ID: UP000000625) from the UniProt Database were collected.[36,37]
2. These IDs were fed into the structure selection algorithm as outlined below and illustrated in the schematic Figure S9. First, X-ray or cryo-EM experimental structures from the Protein Data Bank (PDB) were chosen. When unavailable, the AlphaFold DB model structure was chosen (NMR structures excluded as AlphaFold outperforms NMR in structural accuracy).[38] Other details of the compilation process include the following.
   (a) If multiple X-ray or cryo-EM structures were present, the best one below < 3 Å resolution was selected.
   (b) Structures with more than ten missing residues were excluded and structures with less than six missing residues were shortlisted. If the number of missing residues was between 6 and 10, then this region was modelled using MODELLER.[16]
   (c) Biopython was then used to process these PDB structures to obtain a single chain.[21]

3. Finally, the UniProt IDs that have multiple isoforms (hence no AlphaFold DB model available) were modelled using ColabFold, provided they had more than 16 amino acids.[39]

The compiled ***E. coli* K-12 MG1655 protein structural master database** contained 4382 proteins (out of 4403 total known *E. coli* proteins) (Table S3, schematic in Figure S9). Out of these 1116 had crystal structures. Among these structures, if the PDB dataset (Dataset 2) contained apo and holo versions of the same protein (using a "non-polymer bound ligand query" for each PDB ids), both were considered with an apo/ holo tag. We found 790 apo and 650 holo tagged versions of the RCSB protein structures (Table S3 - Extended Datasets) were added to obtain a total of 4706 structures**.**

**Selecting FAD-binding proteins from the *E. coli* K-12 MG1655 protein structural database**

UniProt IDs of FAD-binding proteins were obtained through a keyword-based search of the UniProt database with the following query: "(taxonomy_id: 83333) AND (keyword: KW-0274)", which stands for *E. coli* (strain K12) and FAD ligand, respectively. Of the 94 output IDs, 70 IDs belong to the *E. coli* K-12 MG1655 strain (Table S6). Upon mapping this to our compiled database, we obtain 24 experimental structures (11 holo- and 13 apo-), and 46 AlphaFold DB models. Since 4 proteins have both apo and holo structures, we obtain 74 FAD-binding protein structures.

**Docking protocol**

Blind docking was performed with DiffDock-L, a diffusion generative model on the *E.coli* K-12 MG1655 protein structural master database.[40] DiffDock-L has demonstrated a 43.0% top-1 success rate (RMSD < 2 Å) on the PDBBind benchmark, significantly outperforming traditional docking methods (smina, GNINA, and P2Rank) as well as other deep learning methods (EquiBind and TANKBind).[41–45] The 3-D structures of FAD, FCD, FGD, and FUD were prepared using OpenBabel after converting their ChemDraw representations into SMILES format and docked with the compiled structural database using batch size of two.[46] Affinity (docking scores) calculations were performed using GNINA with the following parameters: local-only and minimize. Energy minimization was performed with a padding of 2 Å.[42] The relative binding affinity of the FAD analogue was calculated as ΔA (FAD analogue) = A (FAD analogue) – A (FAD), where A denotes the minimum binding affinity among all docking poses for the respective protein and analogue.

DiffDock-L was first validated on eleven holo-FAD *E. coli* proteins having bound FAD ligand obtained from the PDB database. RMSD between the docked and experimental binding poses, using spyrmsd[47] showed that 9 out of 11 complexes (81.8%) had an RMSD < 2 Å. After energy minimisation using GNINA, 8 out of 11 poses (72.7%) had an RMSD < 2 Å. These results demonstrate that DiffDock-L accurately predicts binding poses in holo-FAD proteins. AlphaFold3 (which uses protein sequence and the Ligand SMILES ID) was used to validate DiffDock-L's performance, and it predicts 10 out of 11 structures (90.9%) with an RMSD < 2 Å (Table S4).[48]

To further validate our docking protocol, apo- and AlphaFold DB structures of FAD binding proteins from the *E. coli* K-12 MG1655 protein structural master database was used. Since none of these contain FAD, structural homologs from the RCSB-PDB database that had

FAD bound were identified through a query-based search. The RCSB-PDB database was queried to obtain all the entries with the same "UniProt Molecule Name", provided they also had a bound FAD.[34,35]

Based on resolution, the top-ranked homolog was selected, and the docked protein-ligand complex was aligned with it using PyMol.[14] Eleven such candidates (5 apo structures and 6 AlphaFold DB structures) were found, and visual inspection of the experimental and docking binding sites revealed that the binding site was the same for all. Images of these aligned structures are shown in Figure S11 B,C**.**

**Validating Docking Protocol on Proteins annotated and not annotated as FAD Binding**

To assess the docking protocol's ability to differentiate between FAD-binding and non-FAD-binding proteins, FAD affinities were plotted on a histogram and summary statistics were computed, which showed that these two groups are significantly different (mean affinity for FAD-binding proteins versus non-FAD-binding proteins is -9 kcal/ mol and -3.8 kcal/ mol, respectively). This validates that our docking protocol in distinguishing between the two classes of proteins (Figure S11A).

### SUPPLEMENTARY FIGURES

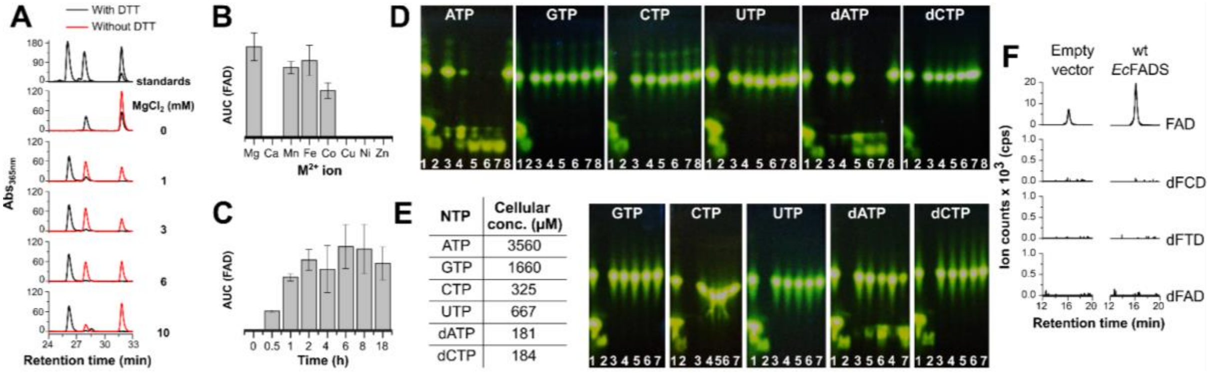

**Figure S1 (related to Figure 2). Activity optimisation of *Ec*FADS. (A)** HPLC chromatogram of *Ec*FADS activity with RF and ATP in the presence of different concentrations of $MgCl_2$ in the presence (black) and absence (red) of DTT. Without DTT, only FMN is formed while with DTT, both FMN and FAD are produced. 3 mM $MgCl_2$ is sufficient to complete the reaction with DTT, while without DTT FMN formation increases until 3 mM $MgCl_2$, and at higher concentrations, the activity decreases. **(B)** Area under the curve (AUC) for FAD peak extracted from HPLC chromatograms of *Ec*FADS activity with various $M^{2+}$ metal ions. Only $Mg^{2+}$, $Mn^{2+}$, $Fe^{2+}$, and $Co^{2+}$ metal ions are accepted by the enzyme in which $Mg^{2+}$ ion is the best for the activity. **(C)** The area under the curve (AUC) for the FAD peak extracted from HPLC chromatograms of *Ec*FADS activity with ATP and RF shows time optimisation for the reaction. From these experiments, we proceeded to do all further experiments with 3 mM $MgCl_2$ in the presence of 10 mM DTT. **(D)** Fluorescent TLC of *Ec*FADS activity with 3.5 mM NTPs and dNTPs. **(E)** TLC of *Ec*FADS activity at the cellular concentration (as given in the table)[2] of NTPs and dNTPs. **(Key for D and E)** Fluorescent TLC of *Ec*FADS activity Lane 1 - RF and FMN standard, Lane 2 - FAD standard, Lane 3 - 1 min reaction, Lane 4 - 10 min reaction, Lane 5- 1 h reaction, Lane 6- 6 h reaction, Lane 7- 15 h reaction. **(F)** EICs of FAD and its deoxy-nucleoside analogues in the strains transformed with empty pET28a(+) and *Ec*FADS in pET28a(+).

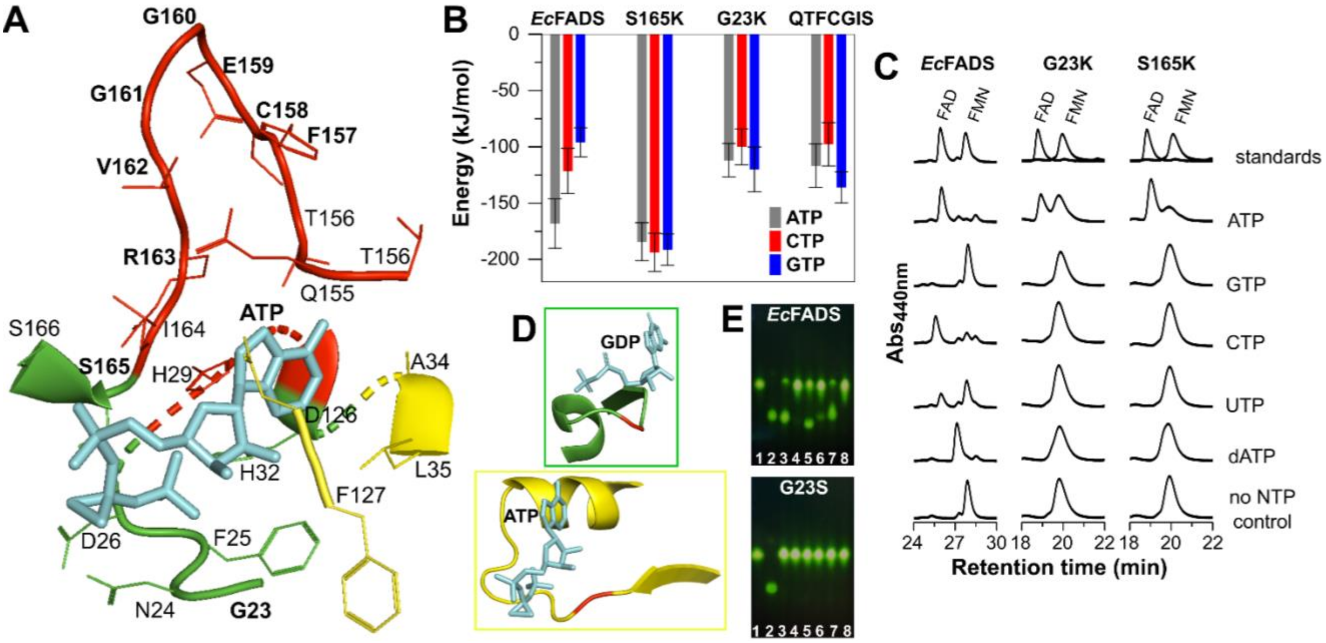

**Figure S2 (related to Figures 3 and 4). Mutations to alter the nucleotide selectivity of *Ec*FADS. (A)** The modelled structure of *Ec*FADS shows the closest residues to ATP where red residues are near the adenine base, green surrounding the phosphates, and yellow are close to the ribose ring. **(B)** Interaction energy for the binding of *Ec*FADS and its mutants with ATP, CTP, and GTP where two residues (G23K and S165K) near the phosphates were mutated with lysine (K), making the negative charge and position of phosphates stable. QTFCGIS mutant was created by shortening the loop (red) around adenine, which shows lower interaction energy for each NTP. **(C)** HPLC chromatogram showing activity assay of *Ec*FADS and its mutants. *Ec*FADS can form FCD, FUD, and dFAD besides FAD. The lysine mutation of G23 and S165 makes the FMNAT domain of *Ec*FADS specific for ATP. **(D)** Glycine (G23) to serine (S) mutation (G23S) for GTP specificity based on a point mutation G105S in FtsZ, a GTPase, in the loop surrounding the phosphates.[49] FtsZ is shown in green, bound to GDP, and the G105 residue is shown in red. *Ec*FADS is in yellow, bound to ATP and G23 residue in red. **(E)** TLC for the activity of *Ec*FADS and G23S mutant with FMN and NTPs. Lane 1- FMN, 2- FAD, 3- reaction with ATP, 4- reaction with GTP, 5- reaction with CTP, 6- reaction with UTP, 7- reaction with dATP, 8- control reaction without NTP.

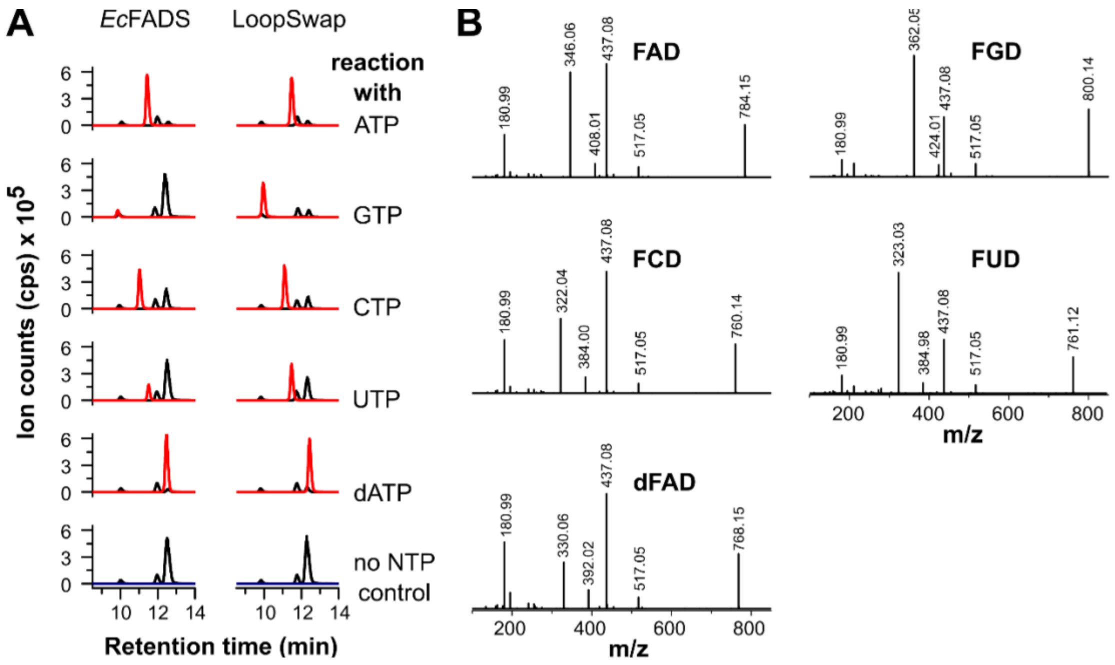

**Figure S3 (related to Figure 3). Activity of LS-*Ec*FADS mutant with NTPs. (A)** EICs for the activity of LS mutant with NTPs compared to WT-*Ec*FADS. FMN was used as starting substrate as shown in black. The EICs for produced FNDs from corresponding NTP have been shown in red. LS mutant accepts all the NTPs and dATP shown and is more efficient. **(B)** Mass fragment analysis of all the FNDs which match with the reported fragment masses.

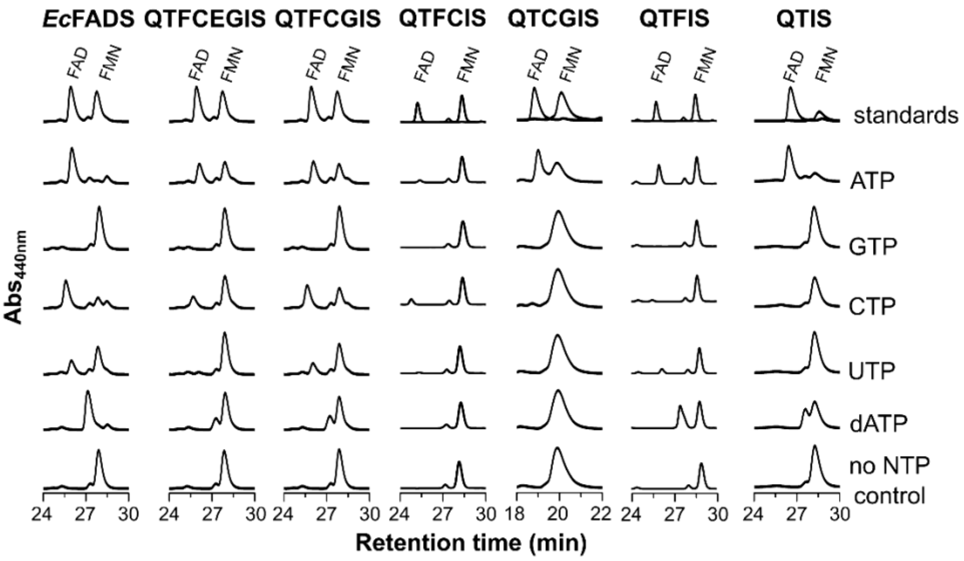

**Figure S4 (related to Figure 4). The activity of WT-*Ec*FADS and its Shortloop mutants with FMN and various NTPs.** HPLC chromatogram showing in vitro activity assay of *Ec*FADS and its mutants. *Ec*FADS can form FCD, FUD, and dFAD besides FAD while on deletion of some residues in the loop region of adenine binding, the activity decreased with each NTP in the case of QTFCEGIS and QTFCGIS and activity further vanished with CTP and UTP for QTFIS and QTIS showing activity with ATP and dATP. However, the QTFCIS and QTCGIS show almost no activity with any NTP except very little for CTP and ATP, respectively.

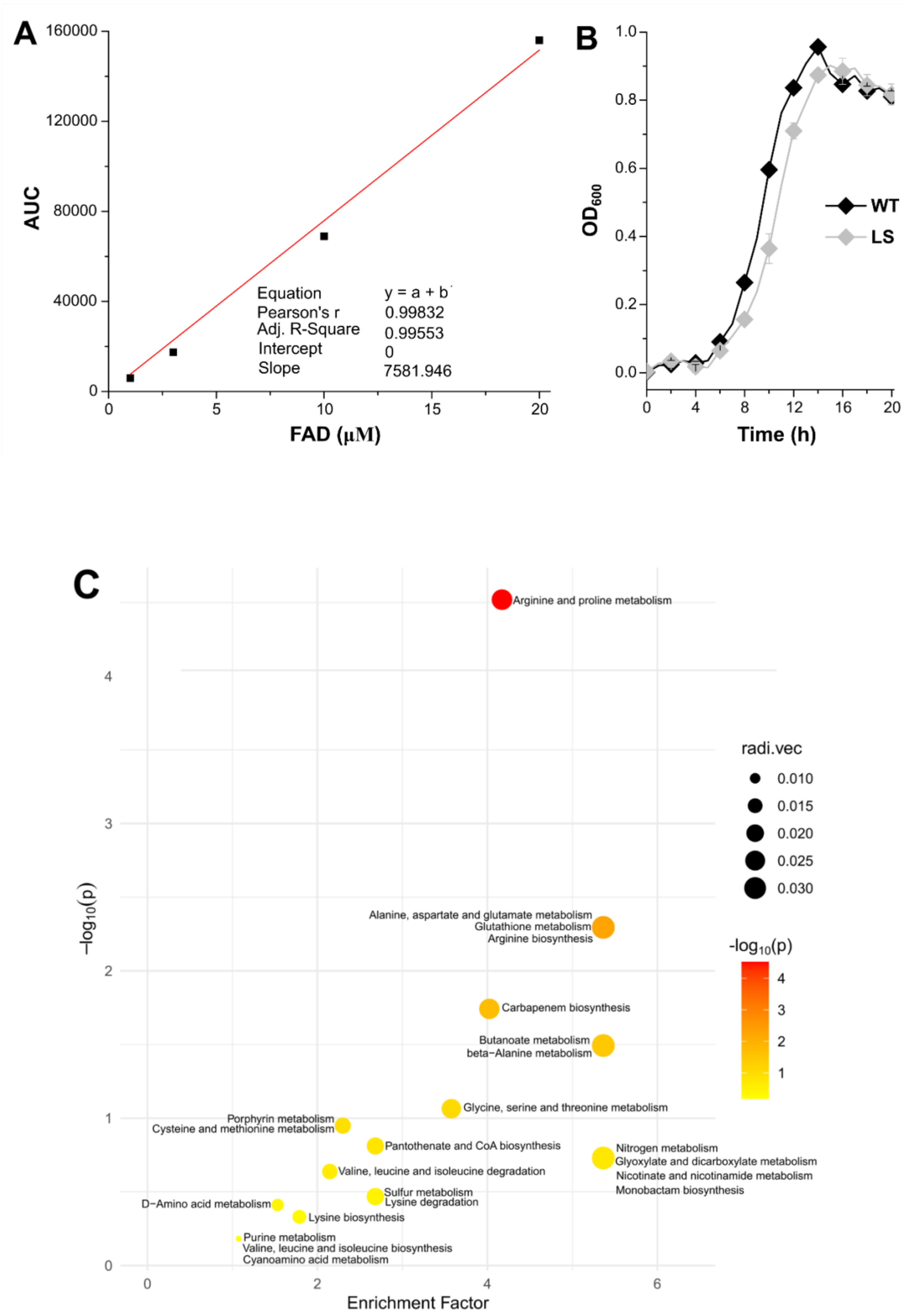

A
AUC
160000
120000
80000
40000
0
FAD (μM)
0
5
10
15
20
Equation
y = a + b
Pearson's r
0.99832
Adj. R-Square
0.99553
Intercept
0
Slope
7581.946
B
OD600
1.0
0.8
0.6
0.4
0.2
0.0
Time (h)
0
4
8
12
16
20
WT
LS
C
Arginine and proline metabolism
Alanine, aspartate and glutamate metabolism
Glutathione metabolism
Arginine biosynthesis
Carbapenem biosynthesis
Butanoate metabolism
beta−Alanine metabolism
Glycine, serine and threonine metabolism
Porphyrin metabolism
Cysteine and methionine metabolism
Pantothenate and CoA biosynthesis
Nitrogen metabolism
Glyoxylate and dicarboxylate metabolism
Nicotinate and nicotinamide metabolism
Monobactam biosynthesis
Valine, leucine and isoleucine degradation
Sulfur metabolism
Lysine degradation
D−Amino acid metabolism
Lysine biosynthesis
Purine metabolism
Valine, leucine and isoleucine biosynthesis
Cyanoamino acid metabolism
−log10(p)
Enrichment Factor
radi.vec
0.010
0.015
0.020
0.025
0.030
-log10(p)

D

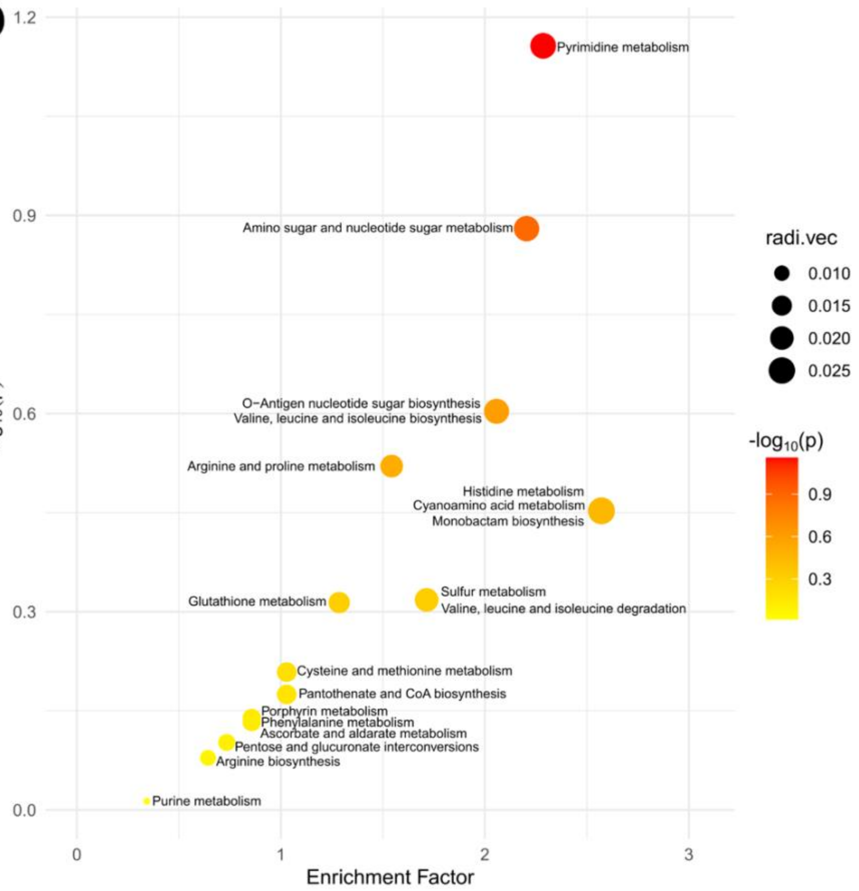
1.2
0.9
0.6
0.3
0.0
−log10(p)
0
1
2
3
Enrichment Factor
Pyrimidine metabolism
Amino sugar and nucleotide sugar metabolism
O−Antigen nucleotide sugar biosynthesis
Valine, leucine and isoleucine biosynthesis
Arginine and proline metabolism
Histidine metabolism
Cyanoamino acid metabolism
Monobactam biosynthesis
Glutathione metabolism
Sulfur metabolism
Valine, leucine and isoleucine degradation
Cysteine and methionine metabolism
Pantothenate and CoA biosynthesis
Porphyrin metabolism
Phenylalanine metabolism
Ascorbate and aldarate metabolism
Pentose and glucuronate interconversions
Arginine biosynthesis
Purine metabolism
radi.vec
0.010
0.015
0.020
0.025
-log10(p)
0.9
0.6
0.3

E

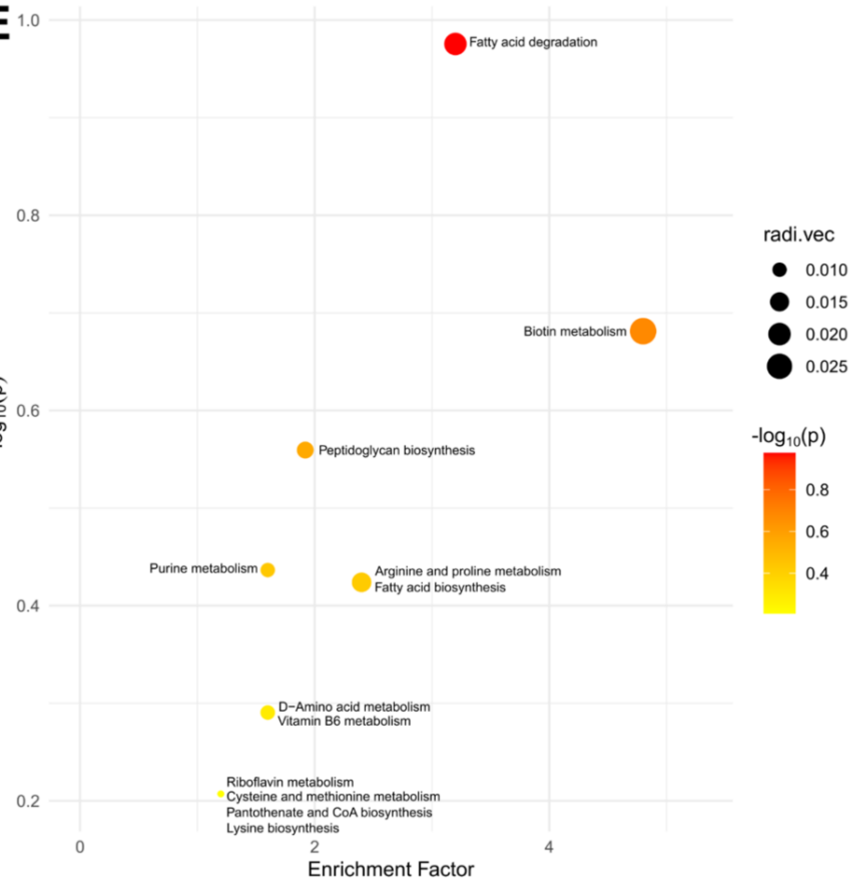
1.0
0.8
0.6
0.4
0.2
−log10(p)
0
2
4
Enrichment Factor
Fatty acid degradation
Biotin metabolism
Peptidoglycan biosynthesis
Purine metabolism
Arginine and proline metabolism
Fatty acid biosynthesis
D−Amino acid metabolism
Vitamin B6 metabolism
Riboflavin metabolism
Cysteine and methionine metabolism
Pantothenate and CoA biosynthesis
Lysine biosynthesis
radi.vec
0.010
0.015
0.020
0.025
-log10(p)
0.8
0.6
0.4

**Figure S5 (Related to Figure 5): (A) LC-MS standard curve of FAD.** This was used to derive the quantity of FAD being produced in the *E. coli* WT and LS strains. **(B)** Growth curve of WT and LS mutant strains in minimal M9 media. **(C and D)** Bubble plot showing the significantly enriched metabolic pathways predicted by Mummichog analysis in **(C)** positive ionization mode and **(D)** negative ionization mode using MetaboAnalyst 6.0.[11,12] **(E)** Bubble plot showing the significantly downriched metabolic pathways predicted by Mummichog analysis in negative ionization mode using MetaboAnalyst 6.0.

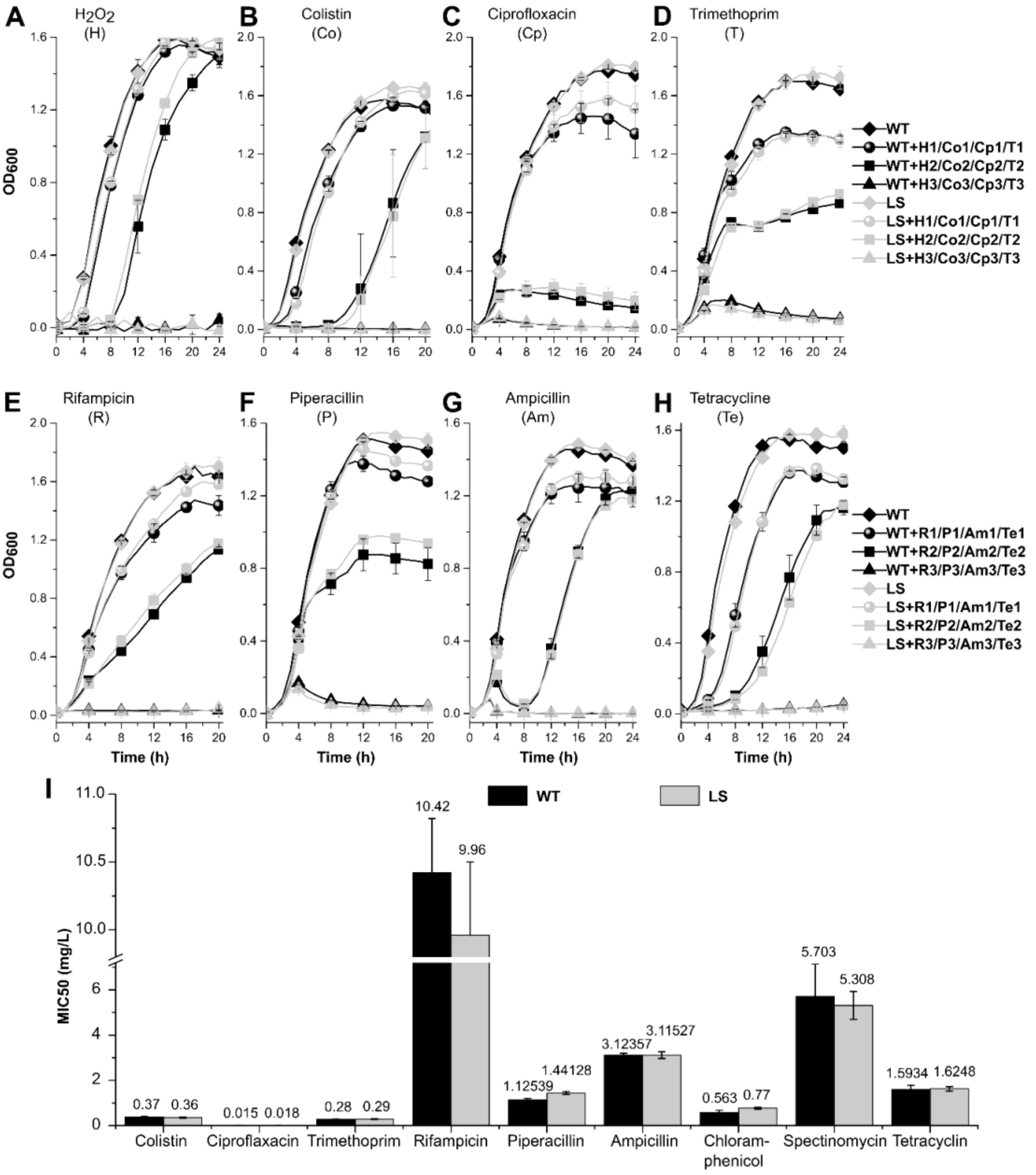

**Figure S6 (related to Figure 6). LS mutant strain against the stress of various antibiotics.** Growth curve of MG1655 WT and LS mutant strain in the presence of **(A)** $H_2O_2$ (H) **(B)** Colistin (Co) **(C)** Ciprofloxacin (Cp) **(D)** Trimethoprim (T) **(E)** Rifampicin (R) **(F)** Piperacillin (P) **(G)** Ampicillin (Am) **(H)** Tetracycline (Te). WT- MG1655 wild-type strain, LS- *Loopswap* mutant strain, H- $H_2O_2$ (H1=1, H2= 2, H3= 5 mM), Co- colistin (Co1= 0.16, Co2= 0.31, Co3= 1.25 μg/mL), Cp- Ciprofloxacin (Cp1= 0.0078, Cp2= 0.0312, Cp3= 0.125 μg/mL), T- Trimethoprim (T1= 0.16, T2= 0.31, T3= 5 μg/mL), R- Rifampicin (R1= 5, R2= 10, R3= 40 μg/mL), P- Piperacillin (P1= 0.62, P2= 1.25, P3= 5 μg/mL), Am- Ampicillin (Am1= 1.25, Am2= 2.5, Am3= 10 μg/mL), Te- Tetracycline (Te1= 0.62, Te2= 1.25, Te3= 5 μg/mL). **(I)** MIC50 values of various antibiotics for WT and LS mutant strains.

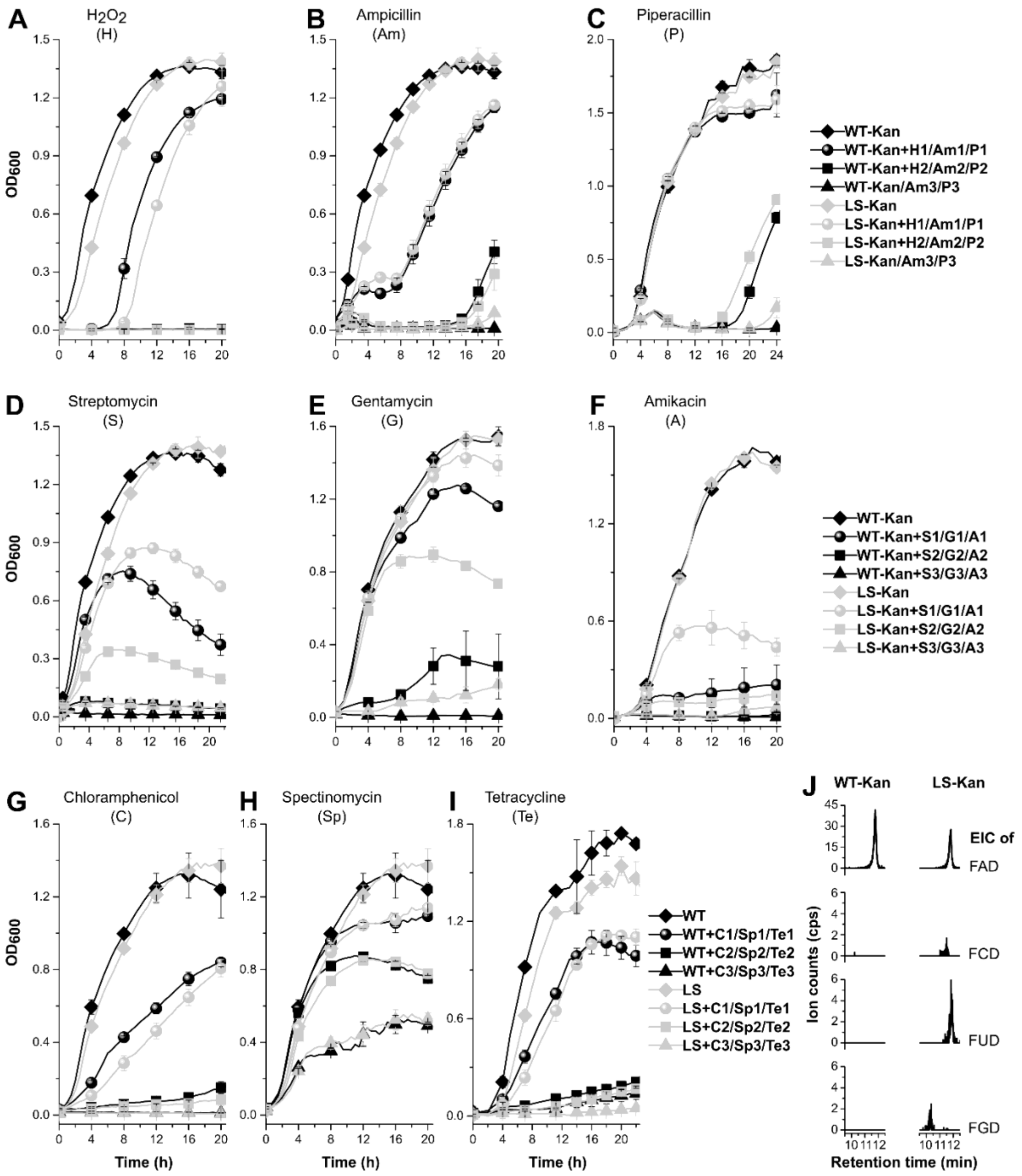

**Figure S7 (related to Figure 5). LS-Kan mutant strain against the stress of various antibiotics and $H_2O_2$.** Growth curve of MG1655 WT-Kan and LS-Kan mutant strain in presence of **(A)** $H_2O_2$ (H) **(B)** Ampicillin (Am) (Cp) **(C)** Piperacillin (P) **(D)** Streptomycin (S) **(E)** Gentamycin (G) **(F)** Amikacin (A) **(G)** Chloramphenicol (C) **(H)** Spectinomycin (Sp) **(I)** Tetracycline (Te). H- $H_2O_2$ (H1=2, H2= 5 mM), Am- Ampicillin (Am1= 2, Am2= 5, Am3= 10 µg/mL), P- Piperacillin (P1= 1, P2= 5, P3= 10 µg/mL), S- Streptomycin (S1= 5, S2= 10, S3= 15 µg/mL), G- Gentamycin (G1= 0.5, G2= 2, G3= 5 µg/mL), A- Amikacin (A1= 5, A2= 10,

A3= 15 μg/mL), C- Chloramphenicol (C1= 1, C2= 2, C3= 6 μg/mL), Sp- Spectinomycin (Sp1= 2, Sp2= 4, Sp3= 10 μg/mL), Te- Tetracycline (Te1= 1, Te2= 2, Te3= 5 μg/mL). **(J)** EICs of FAD and its analogues in the WT-Kan and LS-Kan strains.

| | Q | T | F | C | E | G | G | V | R | I | S | S |
|---|---|---|---|---|---|---|---|---|---|---|---|---|
| A | 13.7 | 3.7 | 0.0 | 13.8 | 2.7 | 8.4 | 2.8 | 8.8 | 0.0 | 8.1 | 0.0 | 0.0 |
| C | 0.0 | 0.1 | 0.4 | 0.8 | 0.4 | 0.1 | 0.1 | 0.2 | 0.0 | 0.0 | 0.0 | 0.0 |
| D | 4.1 | 5.4 | 0.0 | 3.8 | 10.6 | 24.5 | 5.0 | 0.7 | 0.0 | 0.0 | 0.0 | 0.0 |
| E | 3.2 | 1.3 | 0.0 | 26.5 | 9.4 | 7.9 | 2.4 | 25.9 | 0.0 | 0.0 | 0.0 | 0.0 |
| F | 0.0 | 0.0 | 5.6 | 0.2 | 3.1 | 0.0 | 0.0 | 0.0 | 0.0 | 0.1 | 0.0 | 0.0 |
| G | 9.3 | 1.2 | 0.0 | 0.2 | 1.7 | 15.9 | 78.5 | 0.9 | 0.0 | 0.0 | 0.0 | 0.0 |
| H | 8.1 | 0.0 | 0.6 | 0.1 | 9.5 | 19.3 | 0.5 | 4.1 | 0.0 | 0.0 | 0.0 | 0.0 |
| I | 0.2 | 0.0 | 9.5 | 1.7 | 5.7 | 0.1 | 0.0 | 6.6 | 0.0 | 39.2 | 0.0 | 0.0 |
| K | 0.3 | 0.6 | 0.0 | 1.1 | 0.3 | 1.4 | 0.3 | 1.6 | 0.0 | 0.0 | 0.0 | 0.0 |
| L | 1.9 | 0.0 | 6.1 | 10.3 | 10.0 | 0.3 | 0.0 | 18.9 | 0.0 | 0.1 | 0.0 | 0.0 |
| M | 4.2 | 0.1 | 0.1 | 8.7 | 0.9 | 0.1 | 0.0 | 1.2 | 0.0 | 0.0 | 0.0 | 0.0 |
| N | 12.2 | 0.7 | 0.0 | 1.5 | 6.8 | 7.2 | 4.1 | 0.4 | 0.0 | 0.0 | 0.1 | 0.0 |
| P | 29.5 | 1.8 | 0.0 | 0.1 | 0.1 | 0.2 | 0.2 | 0.0 | 0.0 | 0.0 | 0.0 | 0.0 |
| Q | 6.4 | 1.6 | 0.3 | 3.1 | 2.5 | 4.2 | 1.3 | 7.6 | 0.0 | 0.0 | 0.0 | 0.0 |
| R | 2.2 | 0.1 | 0.0 | 1.5 | 1.4 | 1.4 | 0.3 | 1.7 | 99.9 | 0.0 | 0.0 | 0.0 |
| S | 3.2 | 41.5 | 0.0 | 8.6 | 1.3 | 8.2 | 3.8 | 0.9 | 0.0 | 0.2 | 99.9 | 100.0 |
| T | 1.3 | 41.7 | 0.1 | 13.6 | 0.2 | 0.5 | 0.6 | 8.7 | 0.0 | 0.4 | 0.0 | 0.0 |
| V | 0.1 | 0.2 | 55.2 | 4.2 | 32.7 | 0.2 | 0.0 | 11.7 | 0.0 | 51.8 | 0.0 | 0.0 |
| W | 0.0 | 0.0 | 0.0 | 0.0 | 0.4 | 0.0 | 0.0 | 0.0 | 0.0 | 0.0 | 0.0 | 0.0 |
| Y | 0.1 | 0.0 | 22.1 | 0.1 | 0.4 | 0.0 | 0.0 | 0.0 | 0.0 | 0.1 | 0.0 | 0.0 |
| X (Gap) | 0.0 | 0.0 | 0.0 | 0.1 | 0.0 | 0.1 | 0.1 | 0.1 | 0.0 | 0.0 | 0.0 | 0.0 |

**Figure S8. Position specific substitution matrix (PSSM) of the *Ec*FADS loop sequence.** Percentage frequency of each amino acid of the *Ec*FADS FMNAT loop sequence at selected positions in a multiple sequence alignment using 5000 FAD synthetase protein homologues. The rows correspond to the single letter codes of the 20 standard amino acids (plus gaps), and columns represent specific residue positions in the *Ec*FADS FMNAT loop sequence. Each cell contains the percentage frequency with which a given amino acid appears at that position across homologous sequences. Cells with higher values are shaded more intensely, highlighting conserved or strongly preferred residues.

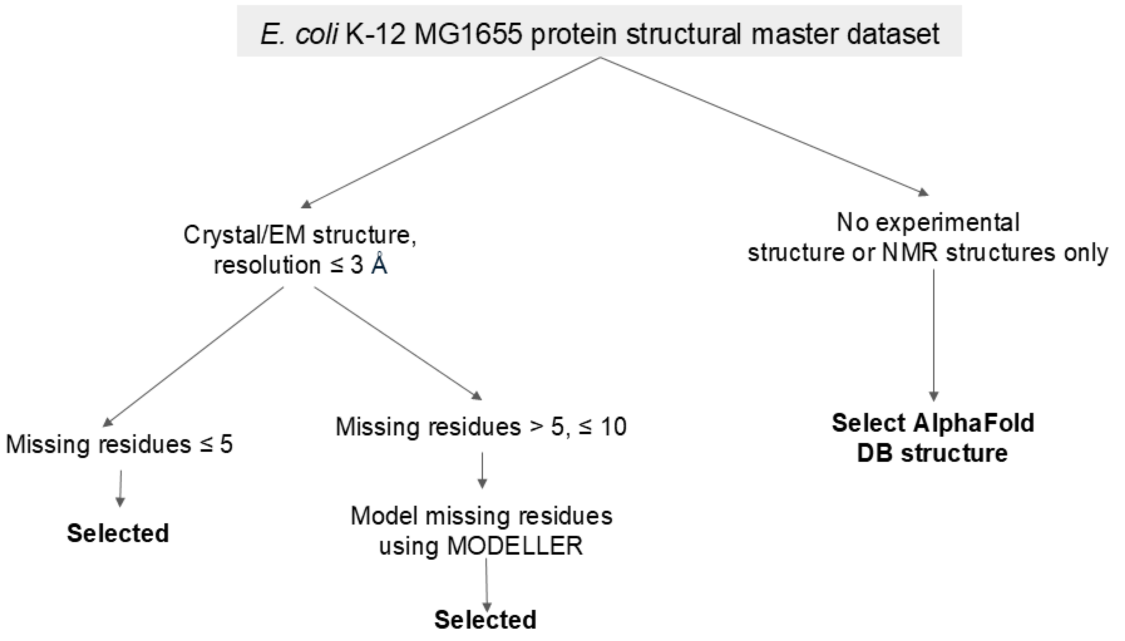

**Figure S9 (Related to Figure 6).** Schematic of the curation of the *E. coli* K-12 MG1655 protein structural master database for docking.

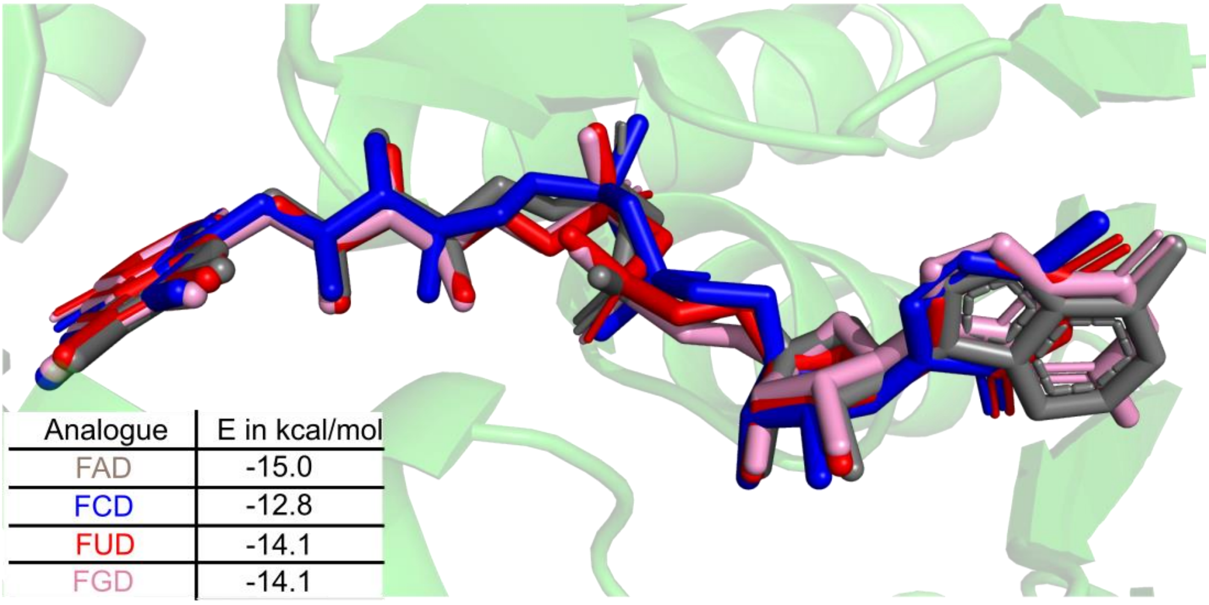

**Figure S10 (Related to Figure 6).** ) Docked structures of FCD, FGD and FUD with the crystal structure of glutathione reductase complexed with FAD (PDB ID: 1GES). The inset table shows the binding affinities calculated by GNINA for each complex.

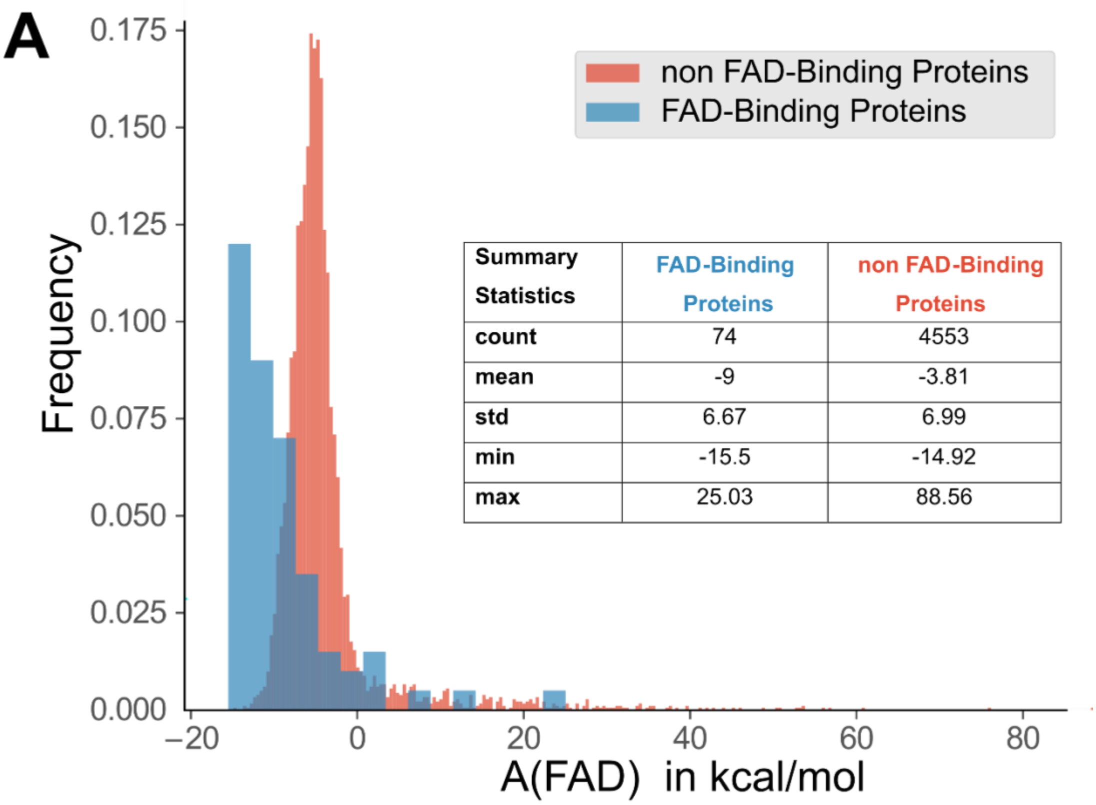

| Summary Statistics | FAD-Binding Proteins | non FAD-Binding Proteins |
|---|---|---|
| count | 74 | 4553 |
| mean | -9 | -3.81 |
| std | 6.67 | 6.99 |
| min | -15.5 | -14.92 |
| max | 25.03 | 88.56 |

**Figure S11 (Related to Figure 6). (A).** Histogram and associated summary statistics (see inset) of FAD affinity for FAD-Binding and non-FAD-Binding proteins.

**B**

| Query UniProt ID | Similar Protein PDB ID | Image of the aligned structures |
|---|---|---|
| P0A6U3 *tRNA uridine 5-carboxymethylaminomethyl modification enzyme MnmG* | 2ZXI | 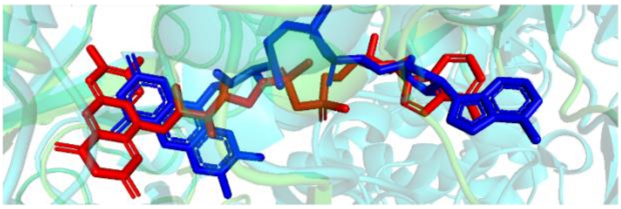 |
| P0A9P0 *Dihydrolipoyl dehydrogenase* | 5U8U | 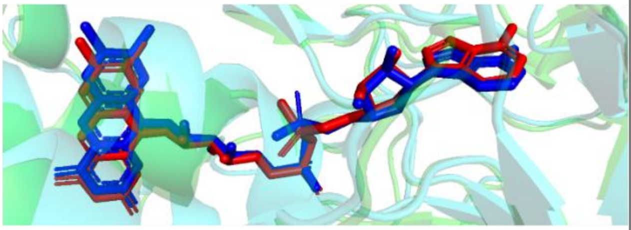 |
| P10902 *L-aspartate oxidase* | 1KNR | 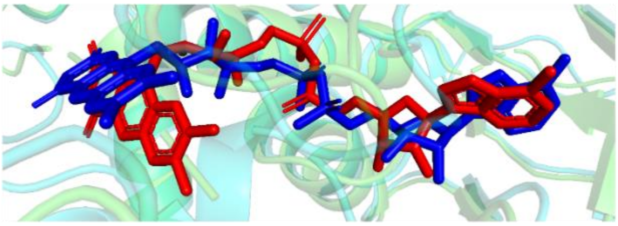 |
| P77212 *Probable pyridine nucleotide-disulfide oxidoreductase* | 6KYY | 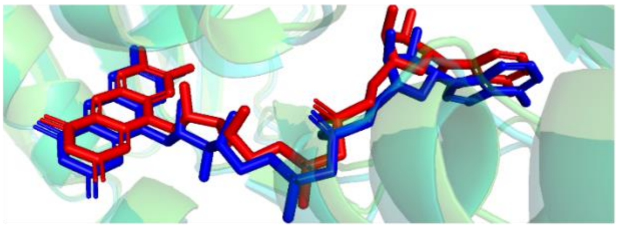 |
| P0A9P4 *Thioredoxin reductase* | 6GNA | 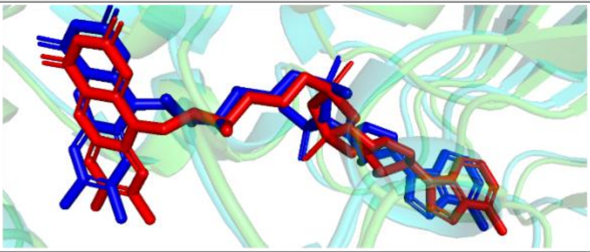 |

**Figure S11 (Related to Figure 6). (B). DiffDock-L predicted top binding pose with Apo FAD-Binding proteins aligned with similar experimental ligand-bound structures.** The red structure is the docked FAD, while the blue structure is the experimentally known pose. The protein structure shown in green is the apo structure, while the one in cyan is the experimental protein structure.

**C**

| Query UniProt ID | Similar Protein PDB ID | Image of the aligned structures |
|---|---|---|
| P00393 *Type II NADH:quinone oxidoreductase* | 5NA1 | 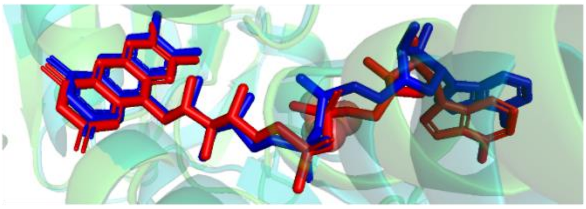 |
| P0AB85 *FAD:protein FMN transferase* | 4IFX | 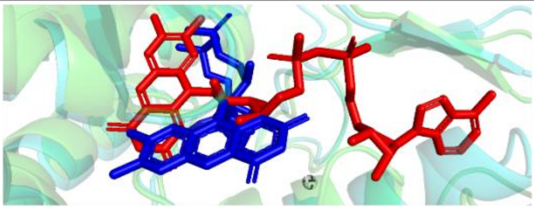 |
| P0AEY5 *NADPH:quinone oxidoreductase MdaB* | 2B3D | 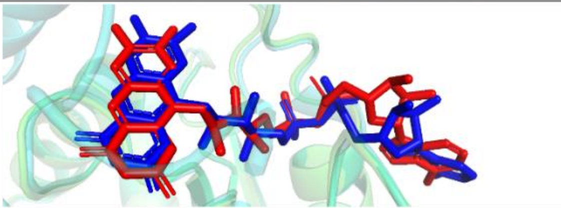 |
| P38038 *Sulfite reductase [NADPH] flavoprotein alpha-component* | 1DDG | 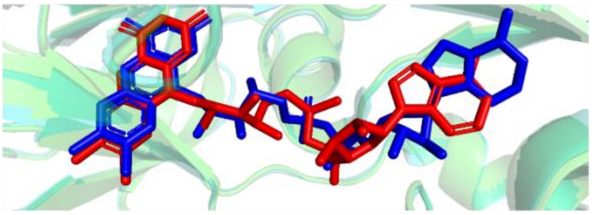 |
| P40874 *N-methyl-L-tryptophan oxidase* | 2UZZ | 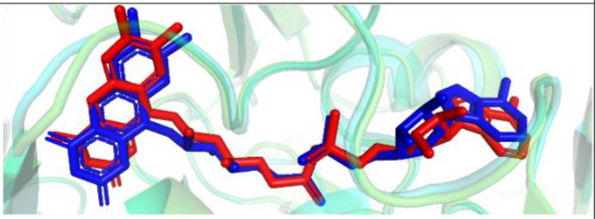 |
| P50466 *Aerotaxis receptor* | 8DIK | 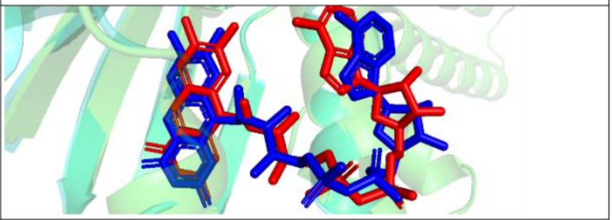 |

**Figure S11 (Related to Figure 6). (C). DiffDock-L predicted top binding pose with AlphaFold2 FAD-Binding proteins aligned with similar experimental ligand-bound structures.** The red structure is the docked FAD, while the blue structure is the experimentally known pose. The protein structure shown in green is the AlphaFold2 structure, while the one in cyan is the experimental protein structure

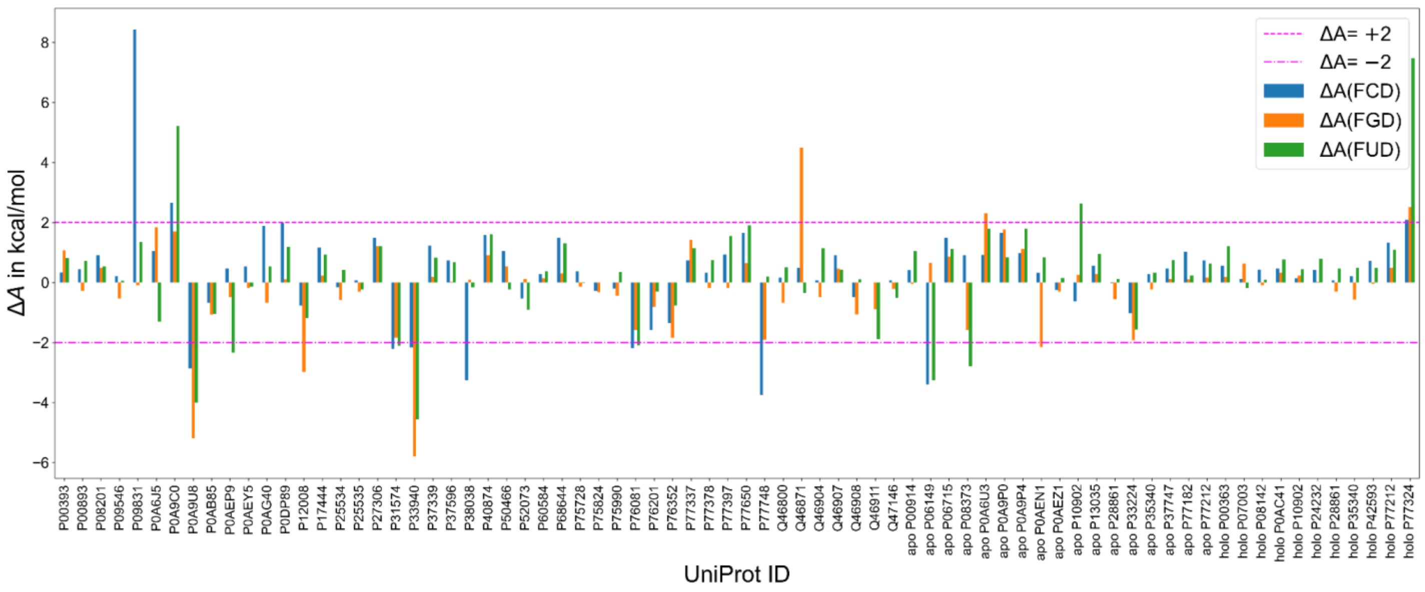

**Figure S12.** (Related to Figure 6). Bar plot of ΔA (FCD, FGD, and FUD) for FAD-binding proteins, where ΔA is defined as the difference between the minimum binding affinity of the analogue and that of FAD [ΔA (FAD analogue) = A (FAD analogue) – A (FAD)].

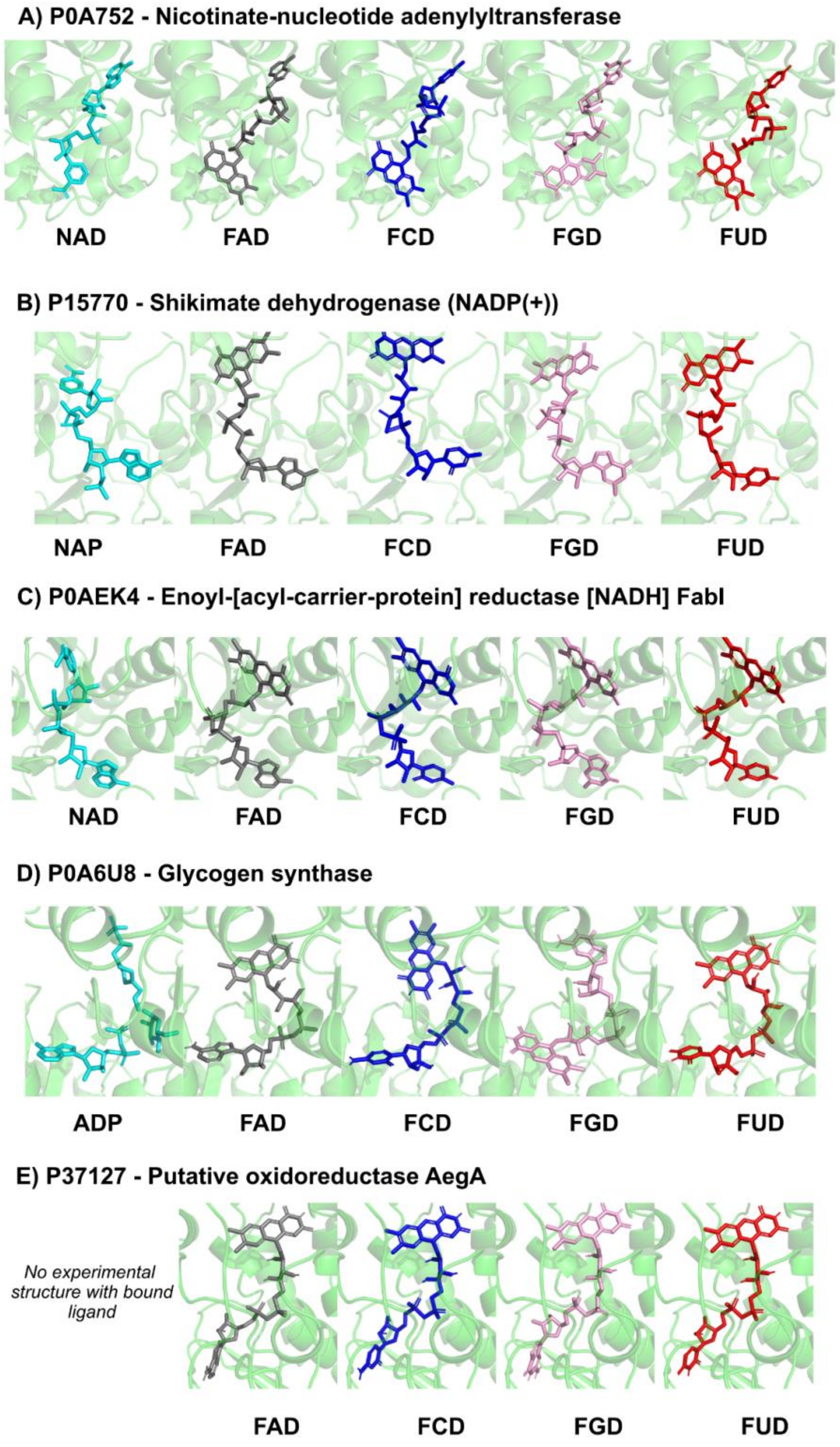

**Figure S13 (Related to Figure 6).** Docked structures of non-FAD binding proteins with FNDs to show better binding toward FAD analogues compared to FAD.

**A) P02931 - Outer membrane porin F**

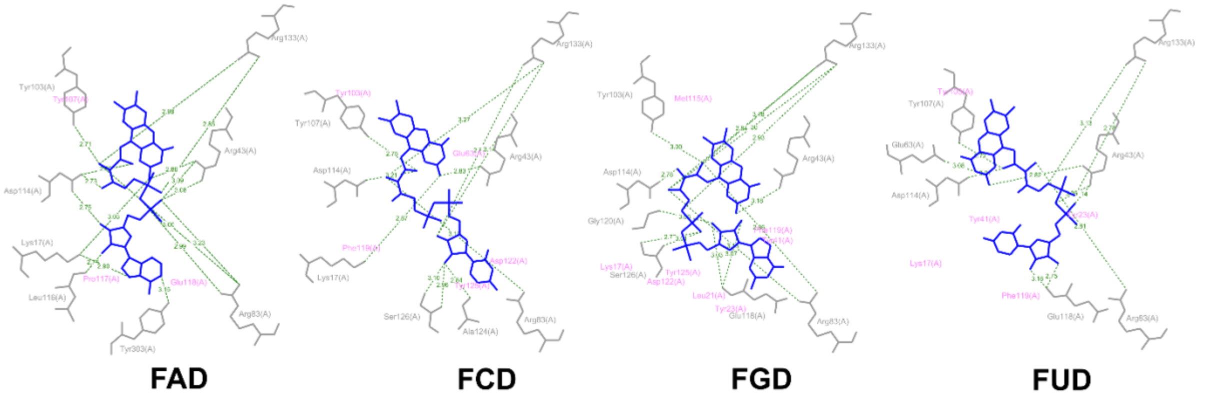

**B) P0AAD2 - Tryptophan-specific transport protein**

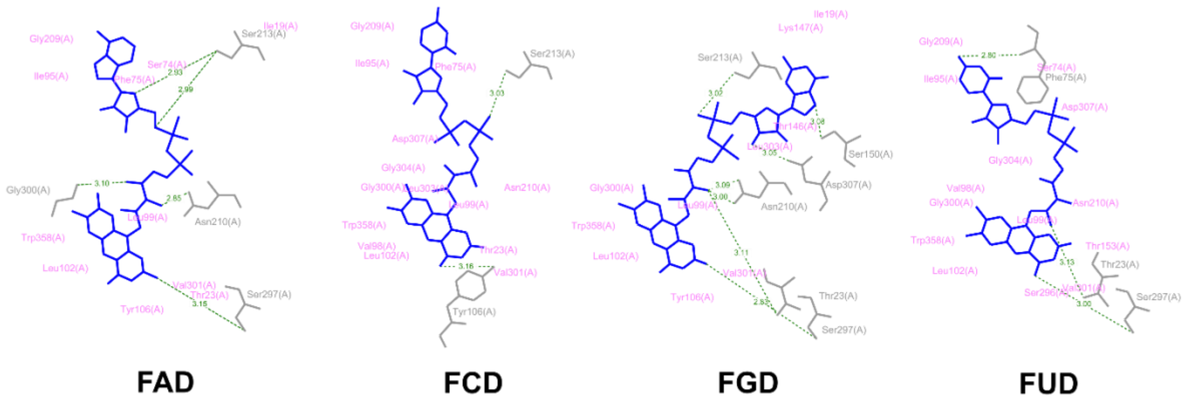

**Figure S14 (Related to Figure 6).** Protein-ligand interaction diagrams for proteins that exhibit stronger binding to FAD analogues (FCD, FGD, and FUD) than to FAD. The selected proteins correspond to genes with (A) the highest upregulation and (B) the highest downregulation based on RNA-seq data.

**Table S1 (related to Figure 3).** List of structural homologs of *Ca*FADS. DALI based search, with Z-score cut-off of 10 was applied.[17] CTP utilizing proteins - glycerol-3-phosphate cytidylyltransferase and choline-phosphate cytidylyltransferase show a loop region around CTP, and glycerol-3-phosphate cytidylyltransferase showed the highest Z-score.

| S. No. | PDB id-Chain | z-score | RMSD | % identity | Structure | Description |
|---|---|---|---|---|---|---|
| 1 | 2x0k-A | 55.2 | 0 | 100 | PDB | RIBOFLAVIN BIOSYNTHESIS PROTEIN RIBF |
| 2 | 3op1-B | 31.5 | 2.2 | 30 | PDB | MACROLIDE-EFFLUX PROTEIN |
| 3 | 1s4m-A | 28.2 | 2.6 | 32 | PDB | RIBOFLAVIN KINASE/FMN ADENYLYLTRANSFERASE |
| 4 | 3q10-A | 14.4 | 5.1 | 16 | PDB | PANTOATE--BETA-ALANINE LIGASE |
| 5 | 3uk2-A | 14.4 | 5.5 | 17 | PDB | PANTOTHENATE SYNTHETASE |
| 6 | 1n06-B | 14.2 | 2.3 | 22 | PDB | PUTATIVE RIBOFLAVIN KINASE |
| 7 | 3inn-C | 14.2 | 5.3 | 12 | PDB | PANTOTHENATE SYNTHETASE |
| 8 | 3ag6-A | 14 | 5.1 | 14 | PDB | PANTOTHENATE SYNTHETASE |
| 9 | 5ucr-A | 13.9 | 6.4 | 18 | PDB | PANTOTHENATE SYNTHETASE |
| 10 | 1iho-A | 13.7 | 4.3 | 15 | PDB | PANTOATE--BETA-ALANINE LIGASE |
| 11 | 1q9s-A | 13.5 | 2.8 | 27 | PDB | HYPOTHETICAL PROTEIN FLJ11149 |
| 12 | 2ejc-A | 13.4 | 5.5 | 14 | PDB | PANTOATE--BETA-ALANINE LIGASE |
| 13 | 7tot-A | 13.4 | 3.8 | 14 | PDB | PANTOTHENATE SYNTHETASE |
| 14 | 3mxt-A | 13.3 | 5.3 | 16 | PDB | PANTOTHENATE SYNTHETASE |
| 15 | 3mue-A | 13.3 | 3.8 | 15 | PDB | PANTOTHENATE SYNTHETASE |
| 16 | 3n8h-A | 13.2 | 8.2 | 14 | PDB | PANTOTHENATE SYNTHETASE |
| 17 | 3q10-D | 13 | 3 | 16 | PDB | PANTOATE--BETA-ALANINE LIGASE |
| **18** | **1coz-A** | **12.7** | **2.5** | **16** | **PDB** | **PROTEIN (GLYCEROL-3-PHOSPHATE)** |
| 19 | 1ufv-A | 12.5 | 3.7 | 16 | PDB | PANTOATE-BETA-ALANINE LIGASE |
| 20 | 3bnw-A | 11.9 | 2.8 | 29 | PDB | RIBOFLAVIN KINASE, PUTATIVE |
| 21 | 3ivc-A | 11.8 | 3.7 | 16 | PDB | PANTOTHENATE SYNTHETASE |
| **22** | **2b7l-A** | **11.3** | **2.7** | **17** | **PDB** | **GLYCEROL-3-PHOSPHATE CYTIDYLYLTRANSFERASE** |

| | | | | | | |
|---|---|---|---|---|---|---|
| 23 | 1v47-A | 11.1 | 2.9 | 15 | PDB | ATP SULFURYLASE |
| 24 | 3k9w-A | 11 | 3.5 | 18 | PDB | PHOSPHOPANTETHEINE ADENYLYLTRANSFERASE |
| 25 | 7wgj-A | 11 | 3.3 | 14 | PDB | PHOSPHOPANTETHEINE ADENYLYLTRANSFERASE |
| 26 | 3otw-A | 10.9 | 3.3 | 19 | PDB | PHOSPHOPANTETHEINE ADENYLYLTRANSFERASE |
| 27 | 3l93-A | 10.9 | 3.2 | 19 | PDB | PHOSPHOPANTETHEINE ADENYLYLTRANSFERASE |
| 28 | 8i8j-B | 10.8 | 3.1 | 15 | PDB | PHOSPHOPANTETHEINE ADENYLYLTRANSFERASE |
| 29 | 1o6b-A | 10.7 | 3.7 | 13 | PDB | PHOSPHOPANTETHEINE ADENYLYLTRANSFERASE |
| 30 | 5h7x-E | 10.6 | 3.5 | 14 | PDB | PHOSPHOPANTETHEINE ADENYLYLTRANSFERASE |
| **31** | **4mvd-B** | **10.5** | **4.2** | **11** | **PDB** | **CHOLINE-PHOSPHATE CYTIDYLYLTRANSFERASE** |
| 32 | 3glv-A | 10.5 | 3.3 | 15 | PDB | LIPOPOLYSACCHARIDE CORE BIOSYNTHESIS PROTEIN |
| 33 | 3nbk-D | 10.4 | 3.4 | 14 | PDB | PHOSPHOPANTETHEINE ADENYLYLTRANSFERASE |
| 34 | 4nah-B | 10.4 | 3.3 | 13 | PDB | PHOSPHOPANTETHEINE ADENYLYLTRANSFERASE |
| 35 | 5o08-A | 10.3 | 3.5 | 14 | PDB | PHOSPHOPANTETHEINE ADENYLYLTRANSFERASE |
| 36 | 5x3d-A | 10.3 | 3.3 | 17 | PDB | PHOSPHOENOLPYRUVATE PHOSPHOMUTASE; |
| 37 | 3nd5-A | 10.2 | 3.2 | 16 | PDB | PHOSPHOPANTETHEINE ADENYLYLTRANSFERASE |
| **38** | **4zcs-A** | **10.2** | **3.9** | **13** | **PDB** | **CHOLINE-PHOSPHATE CYTIDYLYLTRANSFERASE** |
| 39 | 1vlh-B | 10.2 | 3.6 | 16 | PDB | PHOSPHOPANTETHEINE ADENYLYLTRANSFERASE |
| 40 | 5jbn-A | 10.2 | 3.5 | 18 | PDB | PHOSPHOPANTETHEINE ADENYLYLTRANSFERASE |
| 41 | 5ts2-A | 10.2 | 3.5 | 18 | PDB | PHOSPHOPANTETHEINE ADENYLYLTRANSFERASE |
| 42 | 1od6-A | 10.1 | 3.3 | 16 | PDB | PHOSPHOPANTETHEINE ADENYLYLTRANSFERASE |
| 43 | 4f3r-B | 10 | 3.1 | 16 | PDB | PHOSPHOPANTETHEINE ADENYLYLTRANSFERASE |
| 44 | 1f9a-A | 10 | 3.1 | 13 | PDB | HYPOTHETICAL PROTEIN MJ0541 |

**Table S2 (Related to Figures 3,4 and 5).** The list of 80 PDB structures from the PDB database bound to ATP, ADP, AMP, CoA, FAD, NAD, SAM, and their derivatives. [18]

| PDB ID | Ligand | Loop sequence around the nucleobase | N1 H-bonds | N3 H-bonds | N6 Watson-Crick H-bonds | N6 Hoogsteen H-bonds | N7 H-bonds | Name: as listed in PDB database |
|---|---|---|---|---|---|---|---|---|
| 2FT0 | ACO | | HOH/O | | HOH/O | HOH/O | | Crystal structure of TDP-fucosamine acetyltransferase (WecD)- complex with acetyl-CoA |
| 3MQG | ACO | | | | | | HOH/O | crystal structure of the 3-N-acetyl transferase WlbB from Bordetella petrii in complex with acetyl-CoA |
| 5KF9 | ACO | | HOH/O | | | | | X-ray structure of a glucosamine N-Acetyltransferase from Clostridium acetobutylicum in complex with N-acetylglucosamine |
| 1M15 | ADP | | | | S122/O | S282/OG | | Transition state structure of arginine kinase |
| 2C9O | ADP | | V40/N | | V40/O | Y366/OH | | 3D Structure of the human RuvB-like helicase RuvBL1 |
| 2FNA | ADP | | F15/N | K8/NZ | F15/O | HOH/O | HOH/O | Crystal structure of an archaeal aaa+ atpase (sso1545) from sulfolobus solfataricus p2 at 2.00 A resolution |
| **2PYW** | ADP | **RYLEPPH** 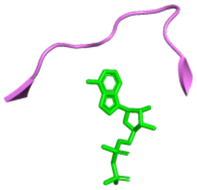 | L119/N | HOH/O | R117/O | | HOH/O | Structure of A. thaliana 5-methylthioribose kinase in complex with ADP and MTR |
| **2XZO** | ADP | **GLPDLN** | | | D470/O | | Q475/NE2 | Upf1 helicase - RNA complex |
| **3D36** | ADP | **TGVGM** | HOH/O | HOH/O | D352/OD2 | HOH/O | HOH/O | How to Switch Off a Histidine Kinase: Crystal Structure of Geobacillus stearothermophilus KinB with the Inhibitor Sda |
| **3EHH** | ADP | **DGTFKG** | HOH/O | K325/N | D320/OD2 | HOH/O | HOH/O | Crystal structure of DesKC-H188V in complex with ADP |
| 3LL3 | ADP | | HOH/O | | HOH/O | HOH/O | | The crystal structure of ligand bound xylulose kinase from Lactobacillus acidophilus |
| 3TTC | ADP | | | | E296/O | HOH/O | HOH/O | Crystal structure of *E. coli* HypF with ADP and carbamoyl phosphate |

| | | | | | | | | |
|---|---|---|---|---|---|---|---|---|
| **3W0O** | ADP | **RAQG** | A95/N | HOH/O | R93/O | HOH/O | HOH/O | Crystal structure of a thermostable mutant of aminoglycoside phosphotransferase APH (4)-Ia, ternary complex with ADP and hygromycin B |
| 4NE2 | ADP | | HOH/O | | | HOH/O | K303/NZ | Pantothenamide-bound Pantothenate Kinase from Klebsiella pneumoniae |
| **4QPM** | ADP | **ELYSYG** | Y869/N | | E867/O | | HOH/O | Structure of Bub1 kinase domain |
| **5FBS** | ADP | **MISPE** | I186/N | | Q184/O | | | Crystal structure of rifampin phosphotransferase RPH-Lm from Listeria monocytogenes in complex with ADP and magnesium |
| 5HE9 | ADP | | | | HOH/O | HOH/O | HOH/O | Bacterial initiation protein in complex with Phage inhibitor protein |
| **5IZ4** | ADP | **ADVG** | V67/N | HOH/O | D66/OD1 | HOH/O | HOH/O | Crystal structure of a putative short-chain dehydrogenase/reductase from Burkholderia xenovorans |
| **5LB3** | ADP | **FGFDSFK** | HOH/O | HOH/O | S28/O | | Q34/NE2 | Crystal structure of human RECQL5 helicase in complex with ADP/Mg. |
| 5LTJ | ADP | | | | | | HOH/O | Crystal structure of the Prp43-ADP-BeF3 complex (in orthorhombic space group) |
| 5LY3 | ADP | | HOH/O | HOH/O | HOH/O/HOH/O | | HOH/O | P. calidifontis crenactin in complex with arcadin-2 C-terminal peptide |
| **12AS** | AMP | **RPDEDRLSPLHSV** | S111/N | | S111/O | E103/OE1 | | ASPARAGINE SYNTHETASE MUTANT C51A, C315A COMPLEXED WITH L-ASPARAGINE AND AMP |
| **1CJA** | AMP | **ELVR** | V165/N | | E163/O | | | ACTIN-FRAGMIN KINASE, CATALYTIC DOMAIN FROM PHYSARUM POLYCEPHALUM |
| 1KHT | AMP | | | | G104/O | T91/OG1 | | Adenylate kinase from Methanococcus voltae |
| **1S68** | AMP | **KIGH** | HOH/O | | | E34/O | I36/N | Structure and Mechanism of RNA Ligase |
| **2GXQ** | AMP | **GLTTP** | HOH/O | | T23/O | | Q28/NE2 | HERA N-terminal domain in complex with AMP, crystal form 1 |
| 3LFR | AMP | | I74/N | HOH/O | I74/O | R96/O | HOH/O | The Crystal Structure of a CBS Domain from a Putative Metal Ion Transporter Bound to AMP from Pseudomonas syringae to 1.55A |

| | | | | | | | | |
|---|---|---|---|---|---|---|---|---|
| **4HG0** | AMP | **RSQMIT** 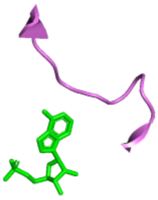 | I80/N | | I80/O | R102/O | | Crystal Structure of magnesium and cobalt efflux protein CorC, Northeast Structural Genomics Consortium (NESG) Target ER40 |
| **5D0N** | AMP | **SGIPRSSPSRNGRL** | | | | | | Crystal structure of maize PDRP bound with AMP |
| 5GMD | AMP | **GLAS** | | | HOH/O | | HOH/O | Crystal structure of Sulfolobus solfataricus diphosphomevalonate decarboxylase in complex with ATP-gamma-S |
| **5NC8** | AMP | **GDAA** | A457/N | H437/N | D456/OD1 | | | Shewanella denitrificans Kef CTD in AMP bound form |
| 1P3D | ANP | | HOH/O | | HOH/O | N295/OD1 | N295/ND2 | Crystal Structure of UDP-N-acetylmuramic acid:L-alanine ligase (MurC) in Complex with UMA and ANP. |
| **3A99** | ANP | **RPEP** | | HOH/O | E121/O | | HOH/O | Structure of PIM-1 kinase crystallized in the presence of P27KIP1 Carboxy-terminal peptide |
| **3C1M** | ANP | **VSGVYTTDP** 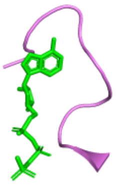 | Y236/N | | Y236/O | HOH/O | | Cyrstal Structure of threonine-sensitive aspartokinase from Methanococcus jannaschii with MgAMP-PNP and L-aspartate |
| 3RC3 | ANP | | HOH/O | HOH/O | | | | Human Mitochondrial Helicase Suv3 |
| 4D2I | ANP | | | | I460/O | | | Crystal structure of the HerA hexameric DNA translocase from Sulfolobus solfataricus bound to AMP-PNP |
| 5FLG | ANP | | V146/N | | V146/O | E31/OE2 | HOH/O | Crystal structure of the 6-carboxyhexanoate-CoA ligase (BioW)from Bacillus subtilis in complex with AMPPNP |
| **5IX1** | ANP | **NGNGM** | | | D67/OD1 | | HOH/O | Crystal structure of mouse Morc3 ATPase-CW cassette in complex with AMPPNP and H3K4me3 peptide |
| **1HP1** | ATP | **PFGN** | N431/ND2 | | | | HOH/O | 5'-NUCLEOTIDASE (OPEN FORM) COMPLEX WITH ATP |
| **1QHH** | ATP | **AHLN** | | Q16/NE2 | | | | STRUCTURE OF DNA HELICASE WITH ADPNP |
| **2A5Y** | ATP | **NVPKQMTCYI** | | | Y131/O | HOH/O | | Structure of a CED-4/CED-9 complex |

| | | | | | | | | |
|---|---|---|---|---|---|---|---|---|
| **3FKQ** | ATP | **APRYEHA** 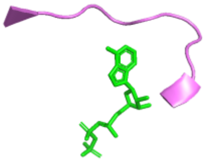 | Y331/N | N307/ND2 | P329/O | N307/OD1 | HOH/O | Crystal structure of NtrC-like two-domain protein (RER070207001320) from Eubacterium rectale at 2.10 A resolution |
| **4CYI** | ATP | **FHPYI** | H340/N | | D338/O | HOH/O | HOH/O | Chaetomium thermophilum Pan3 |
| **4FFL** | ATP | **FDVIK** | V50/N | N73/ND2 | D49/OD1 | | K32/N | PylC in complex with L-lysine |
| 5J1S | ATP | | F70/N | | F70/O | T106/O | HOH/O | TorsinA-LULL1 complex, H. sapiens, bound to VHH-BS2 |
| 2EIS | COA | | | | | HOH/O | | X-ray structure of acyl-CoA hydrolase-like protein, TT1379, from Thermus thermophilus HB8 |
| 4LRT | COA | | HOH/O | | HOH/O | HOH/O | HOH/O | Crystal and solution structures of the bifunctional enzyme (Aldolase/Aldehyde dehydrogenase) from Thermomonospora curvata, reveal a cofactor-binding domain motion during NAD+ and CoA accommodation whithin the shared cofactor-binding site |
| 4PSW | COA | | | | | | | Crystal structure of histone acetyltransferase complex |
| 4U89 | COA | | | HOH/O | | K78/O | HOH/O | 4'-phosphopantetheinyl transferase PptT from Mycobacterium tuberculosis |
| 2CUL | FAD | | A91/N | Q34/N | A91/O | HOH/O | HOH/O | Crystal structure of the GidA-related protein from Thermus thermophilus HB8 |
| 2IPI | FAD | | V203/N | | V203/O | S67/OG | S67/OG | Crystal Structure of Aclacinomycin Oxidoreductase |
| 2QCU | FAD | | A172/N | A34/N | | HOH/O | | Crystal structure of Glycerol-3-phosphate Dehydrogenase from Escherichia coli |
| 3DME | FAD | | L173/N | A35/N | L173/O | HOH/O | HOH/O | Crystal structure of conserved exported protein from Bordetella pertussis. NorthEast Structural Genomics target BeR141 |
| 3EC6 | FAD | | | HOH/O | | | | Crystal structure of the General Stress Protein 26 from Bacillus anthracis str. Sterne |
| 3EWK | FAD | | | | | | | Structure of the redox sensor domain of Methylococcus capsulatus (Bath) MmoS |

| | | | | | | | | |
|---|---|---|---|---|---|---|---|---|
| 3F8D | FAD | | V92/N | E46/N | V92/O | | | Structure of Sulfolobus solfataricus Thioredoxin reductase Mutant C147A |
| 3JQQ | FAD | | | | | HOH/O | | Crystal structure of the H286K mutant of Ferredoxin-NADP+ reductase from Plasmodium falciparum in complex with 2'P-AMP |
| 3NKS | FAD | | V257/N | S35/N | V257/O | | HOH/O | Structure of human protoporphyrinogen IX oxidase |
| 4EQS | FAD | | V81/N | K34/N | V81/O | HOH/O | HOH/O | Crystal structure of the Y419F mutant of Staphylococcus aureus CoADR |
| 4H4R | FAD | | A82/N | D40/N | A82/O | HOH/O | HOH/O | Crystal Structure of Ferredoxin reductase, BphA4 E175C/Q177G mutant (oxidized form) |
| 4X9M | FAD | | V177/N | K34/N | V177/O | | | Oxidized L-alpha-Glycerophosphate Oxidase from Mycoplasma pneumoniae with FAD bound |
| 5JCI | FAD | | I96/N | K40/N | I96/O | HOH/O | HOH/O | Structure and catalytic mechanism of monodehydroascorbate reductase, MDHAR, from Oryza sativa L. japonica |
| 5KF6 | FAD | | | | T393/O | | | Structure of proline utilization A from Sinorhizobium meliloti complexed with L-tetrahydrofuroic acid and NAD+ in space group P21 |
| 1KAE | NAD | | | HOH/O | | | | L-HISTIDINOL DEHYDROGENASE (HISD) STRUCTURE COMPLEXED WITH L-HISTIDINOL (SUBSTRATE), ZINC AND NAD (COFACTOR) |
| 4CPD | NAD | | | | | | | Alcohol dehydrogenase TADH from Thermus sp. ATN1 |
| 5TT5 | NAD | | K290/NZ | HOH/O | E113/OE2 | L114/O | L116/N | Escherichia coli LigA (K115M) in complex with NAD+ |
| 2C0C | NAP | | HOH/O | HOH/O | HOH/O | HOH/O | | Structure of the MGC45594 gene product |
| 2Y5D | NAP | | HOH/O | | | | | Crystal structure of C296A mutant of the box pathway encoded ALDH from Burkholderia xenovorans LB400 |
| 3LZW | NAP | | HOH/O | HOH/O | | | | Crystal structure of ferredoxin-NADP+ |

| | | | | | | | | |
|---|---|---|---|---|---|---|---|---|
| | | | | | | | | oxidoreductase from bacillus subtilis (form I) |
| 4DPL | NAP | | HOH/O | HOH/O | | | | Structure of malonyl-coenzyme A reductase from crenarchaeota in complex with NadP |
| 4R3N | NAP | | HOH/O | | | | | Crystal structure of the ternary complex of sp-ASADH with NADP and 1,2,3-Benzenetricarboxylic acid |
| 5DP2 | NAP | | HOH/O | HOH/O | HOH/O/HOH/O | HOH/O | HOH/O | CurF ER cyclopropanase from curacin A biosynthetic pathway |
| **2HA8** | SAH | **IPQQGIIRSL** 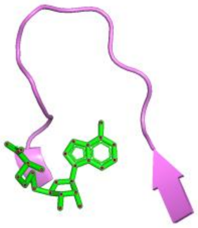 | I149/N | | I149/O | | L158/N | Methyltransferase Domain of Human TAR (HIV-1) RNA binding protein 1 |
| 2JJQ | SAH | | D326/N | S300/N | D326/OD1 | HOH/O | | The crystal structure of Pyrococcus abyssi tRNA (uracil-54, C5)-methyltransferase in complex with S-adenosyl-L-homocysteine |
| 3H2B | SAH | | I150/N | | HOH/O | | HOH/O | Crystal structure of the SAM-dependent methyltransferase cg3271 from Corynebacterium glutamicum in complex with S-adenosyl-L-homocysteine and pyrophosphate. Northeast Structural Genomics Consortium Target CgR113A |
| **3NDC** | SAH | **RASWPDQ** | A193/N | | A193/O | HOH/O/P10/O | HOH/O/HOH/O | Crystal structure of Precorrin-4 C11-methyltransferase from Rhodobacter capsulatus |
| **3TOS** | SAH | **TFTGFPDV** | V167/N | T108/N | D166/OD1 | HOH/O | | Crystal Structure of CalS11, Calicheamicin Methyltransferase |
| 4JWJ | SAH | | L232/N | | HOH/O | R244/O | L246/N | Crystal structure of scTrm10(84)-SAH complex |
| 5CCB | SAH | | V164/N | F136/N | D163/OD1 | | | Crystal structure of human m1A58 methyltransferase in a complex with tRNA3Lys and SAH |
| **5MGZ** | SAH | **GDAQLLAG** | A97/N | L71/N | D96/OD1 | E4/OE1 | | Streptomyces Spheroides NovO (8-demethylnovbiocic acid methyltransferase) with SAH |

| ACO: Acetyl coenzyme A; ADP: Adenosine diphosphate; AMP: Adenosine monophosphate; ATP: Adenosine triphosphate; CoA: Coenzyme A; FAD: Flavin adenine dinucleotide; NAD: Nicotinamide adenine dinucleotide; NAP: Nicotinamide adenine dinucleotide phosphate; SAH: S-adenosylhomocysteine |
|---|

**Table S3 (related to Figure 6)**. Dataset ID and corresponding sources of the protein structures in the ***E. coli*** **K-12 MG1655 protein structural master database.**

| **Dataset ID** | **Number of Proteins** | **Source of the Structures and their Description** |
|---|---|---|
| 1 | 2715 | AlphaFold DB<br>(structures that have only AlphaFold DB structures) |
| 2 | 1116 | RCSB PDB<br>(X-ray or cryo-EM experimental structures satisfying resolution and missing residue cutoffs) |
| 3 | 17 | ColabFold<br>(structures neither present in RCSB PDB nor in AlphaFold DB) |
| 4 | 9 | RCSB PDB, missing residues modelled using MODELLER.<br>(X-ray or cryo-EM experimental structures satisfying resolution cutoff but having between 6 to 10 missing residues.) |
| 6 | 525 | AlphaFold DB<br>(Proteins that have X-ray or cryo-EM experimental structures that do not meet resolution or missing residue cutoff/ proteins that have only NMR structures.) |
| | 4382* | Total number of proteins in the compiled dataset |
| | 4403 | Total number of proteins in *E. coli* K-12 MG1655 |
| **Extended Datasets** | | |
| 9 | 790 | RCSB PDB<br>(X-ray or cryo-EM experimental structures that satisfy resolution and missing residues cutoff and do not have bound ligands) |
| 10 | 650 | RCSB PDB<br>(X-ray or cryo-EM experimental structures that satisfy resolution and missing residues cutoff and have bound ligands) |

*21 proteins with less than 16 amino acids without an experimental or computed structure were not included in the dataset.

**Dataset IDs 5, 7 and 8 were not used in any codes accompanying this manuscript.

**Table S4 (related to Figure 6).** RMSDs of first ranked DiffDock-L, DiffDock-L + GNINA and AlphaFold3 posed with holo *E. coli* MG-1655 proteins against their crystallised FAD structures.

| **Protein Name** | **UniProt ID** | **RMSD (Å) with DiffDock-L docked pose** | **RMSD (Å) with GNINA minimised pose** | **RMSD (Å) with AlphaFold 3 docked pose** |
|---|---|---|---|---|
| Fumarate reductase flavoprotein subunit | P00363 | 0.86 | 0.99 | 0.27 |
| Pyruvate dehydrogenase [ubiquinone] | P07003 | 0.51 | 0.67 | 0.54 |
| Acetolactate synthase isozyme 1 large subunit | P08142 | 1.23 | 1.53 | 0.36 |
| Succinate dehydrogenase flavoprotein subunit | P0AC41 | 0.84 | 1.01 | 0.43 |
| L-aspartate oxidase | P10902 | 0.75 | 0.67 | 0.43 |
| Flavohemoprotein | P24232 | 4.84 | 5.01 | 0.52 |
| Flavodoxin/ferredoxin--NADP reductase | P28861 | 1.83 | 2.33 | 0.27 |
| Alkyl hydroperoxide reductase subunit F | P35340 | 0.96 | 0.81 | 23.89 |
| 2,4-dienoyl-CoA reductase [(2E)-enoyl-CoA-producing] | P42593 | 1.9 | 1.08 | 0.55 |
| Probable pyridine nucleotide-disulfide oxidoreductase RclA | P77212 | 0.74 | 0.67 | 0.36 |
| Aldehyde oxidoreductase FAD-binding subunit PaoB | P77324 | 18.52 | 20.59 | 0.42 |

**Table S5 (related to Figure 6)**. ΔA values of 9 holo FAD-binding proteins with RMSD <2 Å.

| **Sl. no.** | **UniProt ID** | **Protein Name** | **A (FAD) in kcal/mol** | **ΔA (FCD) in kcal/mol** | **ΔA (FGD) in kcal/mol** | **ΔA (FUD) in kcal/mol** |
|---|---|---|---|---|---|---|
| 1 | P28861 | Flavodoxin/ferredoxin--NADP reductase | -10.42 | 1.13 | 0.11 | -0.27 |
| 2 | P07003 | **Pyruvate dehydrogenase [ubiquinone]** | -14.18 | 0.67 | 0.62 | 0.9 |
| 3 | P00363 | Fumarate reductase flavoprotein subunit | -13.76 | -0.65 | -1.04 | -1.04 |
| 4 | P77212 | Probable pyridine nucleotide-disulfide oxidoreductase RclA | -14.17 | 1.55 | 0.49 | 1.26 |
| 5 | P42593 | **2,4-dienoyl-CoA reductase [(2E)-enoyl-CoA-producing]** | -13.34 | 0.52 | 2.07 | 0.33 |
| 6 | P24232 | Flavohemoprotein | -12.95 | 1.26 | -0.65 | 0.12 |
| 7 | P10902 | L-aspartate oxidase | -15.15 | -0.5 | -0.3 | 0.06 |
| 8 | P08142 | Acetolactate synthase isozyme 1 large subunit | -13.6 | 0.46 | 0.77 | -0.06 |
| 9 | P0AC41 | Succinate dehydrogenase flavoprotein subunit | -13.58 | 2.61 | 0.84 | 1.75 |

**Table S6 (related to Figure 6).** UniProt IDs of FAD-binding proteins in *E. coli* K-12 MG1655 strain.

| **UniProt ID** | **UniProt ID** | **UniProt ID** | **UniProt ID** |
|---|---|---|---|
| P00363 | P0AEY5 | P76081 | P31574 |
| P00393 | P0AEZ1 | P77182 | Q46908 |
| P00893 | P10902 | P77212 | Q46911 |
| P00914 | P12008 | P77324 | P0A9U8 |
| P06149 | P13035 | P77748 | P68644 |
| P06715 | P17444 | Q46871 | P76201 |
| P07003 | P24232 | Q47146 | P77337 |
| P08142 | P25534 | P0AB85 | P77378 |
| P08201 | P25535 | P0AG40 | Q46904 |
| P08373 | P27306 | P33940 | Q46907 |
| P09546 | P28861 | P37596 | |
| P09831 | P33224 | P50466 | |
| P0A6J5 | P35340 | P60584 | |
| P0A6U3 | P37339 | P75728 | |
| P0A9C0 | P37747 | P75824 | |
| P0A9P0 | P38038 | P76352 | |
| P0A9P4 | P40874 | P77397 | |
| P0AC41 | P42593 | P77650 | |
| P0AEN1 | P52073 | Q46800 | |
| P0AEP9 | P75990 | P0DP89 | |

**Table S7(related to Figure 6).** Affinities of non-FAD binding proteins that bind stronger to FAD analogues than to FAD based on affinity and confidence scores.

| | **Composition in top confidence scores** | | | | **Composition in top A** | | | | | |
|---|---|---|---|---|---|---|---|---|---|---|
| **UniProt ID** | **FAD** | **FC D** | **FGD** | **FUD** | **FAD** | **FCD** | **FGD** | **FUD** | **Protein name** | **Structure type** |
| P37127 | 1 | 1 | 5 | 3 | 0 | 1 | 2 | 7 | AEGA_ECOLI Putative oxidoreductase AegA | AlphaFold DB |
| P0A6U8 | 1 | 3 | 3 | 3 | 3 | 1 | 4 | 2 | GLGA_ECOLI Glycogen synthase | Apo |
| P0A6U8 | 4 | 3 | 3 | 0 | 4 | 0 | 6 | 0 | GLGA_ECOLI Glycogen synthase | Holo |
| P0A752 | 2 | 2 | 4 | 2 | 4 | 3 | 2 | 1 | NADD_ECOLI Nicotinate-nucleotide adenylyltransferase | Apo |
| P0AEK4 | 3 | 1 | 3 | 3 | 4 | 1 | 2 | 3 | FABI_ECOLI Enoyl-[acyl-carrier-protein] reductase [NADH] FabI | Apo |
| P15770 | 3 | 3 | 2 | 2 | 3 | 0 | 4 | 3 | AROE_ECOLI Shikimate dehydrogenase (NADP (+)) | Apo |

**Table S8 (related to Figure 6).** Affinities of non-FAD-binding proteins that bind FAD analogues more strongly than FAD, with corresponding RNA-seq upregulation or downregulation.

| UniProt ID | Description | Fold Change (LS vs WT) | Regulation | Count (FAD) | Count (FCD) | Count (FGD) | Count (FUD) | FAD Binding Protein | Affinity Range (kcal/mol) |
|---|---|---|---|---|---|---|---|---|---|
| P0AAD2 | MTR_ECOLI Tryptophan-specific transport protein | -48.046032 | down | 3 | 4 | 1 | 2 | FALSE | -9.6 to -8.7 |
| P26459 | APPC_ECOLI Cytochrome bd-II ubiquinol oxidase subunit 1 | -37.346172 | down | 2 | 2 | 1 | 5 | FALSE | -11.7 to -7.9 |
| P33224 | AIDB_ECOLI Putative acyl-CoA dehydrogenase AidB | -15.201208 | down | 2 | 3 | 2 | 3 | TRUE | -10.6 to -7.9 |
| P0AEZ1 | METF_ECOLI 5,10-methylenetetrahydrofolate reductase | -11.880665 | down | 3 | 2 | 4 | 1 | TRUE | -12.8 to -12.4 |
| P08190 | FIMG_ECOLI Protein FimG | 10.029439 | up | 0 | 1 | 5 | 4 | FALSE | -8.8 to -8.2 |
| P08189 | FIMF_ECOLI Protein FimF | 11.222717 | up | 3 | 2 | 1 | 4 | FALSE | -8.6 to -7.7 |
| P39264 | FIMI_ECOLI Fimbrin-like protein FimI | 12.283414 | up | 2 | 4 | 3 | 1 | FALSE | -8.8 to -7.8 |
| P0AEX9 | MALE_ECOLI Maltose/maltodextrin-binding periplasmic protein | 13.884436 | up | 2 | 2 | 3 | 3 | FALSE | -10.5 to -8.7 |
| P0AAG8 | MGLA_ECOLI Galactose/methyl galactoside import ATP-binding protein MglA | 18.850134 | up | 3 | 4 | 2 | 1 | FALSE | -10.1 to -8.7 |
| P02943 | LAMB_ECOLI Maltoporin | 62.38295 | up | 2 | 2 | 4 | 2 | FALSE | -10 to -8.3 |
| P0AE22 | APHA_ECOLI Class B acid phosphatase | 67.57143 | up | 4 | 2 | 1 | 3 | FALSE | -10.2 to -7.9 |
| P02931 | OMPF_ECOLI Outer membrane porin F | 95.65126 | up | 2 | 3 | 4 | 1 | FALSE | -9.8 to -7.9 |

**Table S9: Common hits from RNA sequencing and Metabolomics study**

| List of Upregulated pathways | Glutamate as a hit | Maps to RNASeq data | |
|---|---|---|---|
| Arginine biosynthesis | Yes | | |
| Alanine, aspartate and glutamate metabolism | Yes | | |
| Glutathione metabolism | Yes | Yes | Taurine & Hypotaurine metabolism |
| Carbapenem biosynthesis | Yes | | |
| beta-Alanine metabolism | | | |
| Butanoate metabolism | Yes | | |
| Glycine, serine and threonine metabolism | | | |
| Porphyrin metabolism | Yes | | |
| Monobactam biosynthesis | | | |
| Glyoxylate and dicarboxylate metabolism | Yes | Yes | |
| Nicotinate and nicotinamide metabolism | Yes | | |
| Nitrogen metabolism | Yes | Yes | |
| Valine, leucine and isoleucine degradation | | | |
| Lysine degradation | | | |
| One carbon pool by folate | | | |
| Sulfur metabolism | Yes | Yes | Taurine & Hypotaurine metabolism |
| Valine, leucine and isoleucine biosynthesis | | | |
| Cyanoamino acid metabolism | | | |
| Pyrimidine metabolism | | | |
| Amino sugar and nucleotide sugar metabolism | | | |
| O-Antigen nucleotide sugar biosynthesis | | | |
| Histidine metabolism | | | |
| Phenylalanine metabolism | | | |
| Ascorbate and aldarate metabolism | | | |
| Pentose and glucuronate interconversions | | | |

**Table S10: List of primers used for cloning.**

| # | Primer name | Sequence |
|---|---|---|
| 1 | *Ec*FADS_F1 | 5'-ATGAAGCTGATACGCGGCATAC-3' |
| 2 | *Ec*FADS_R1 | 5'-TTAAGCCGGTTTTGTTAGCCC-3' |
| 3 | *Ec*FADS_F2 | 5'-GTGCCGCGCGGCAGCCATATGAAGCTGATACGCGGCATAC-3' |
| 4 | *Ec*FADS_R2 | 5'-CGACGGAGCTCGAATTCGGATCCTTAAGCCGGTTTTGTTAGCCC-3' |
| 5 | G23K_F | 5'-GCTGACTATTAAAAATTTCGACG-3' |
| 6 | S165K_F | 5'-GCGCATCAAAAGCACCGC-3' |
| 7 | G23S_F | 5'-GCTGACTATTAGCAATTTCGACG-3' |
| 8 | QTFCEGIS_F | 5'-ACGCAAACTTTTTGCGAAGGTATCAGCAGCACCGCC-3' |
| 9 | QTFCGIS_F | 5'-CCAGTACGCAAACTTTTTGCGGTATCAGCAGCACCGCC-3' |
| 10 | QTFCIS_F | 5'-CACCAGTACGCAAACTTTTTGCATCAGCAGCACCGCCG-3' |
| 11 | QTCGIS_F | 5'-TCACCAGTACGCAAACTTGCGGCATCAGCAGCACCG-3' |
| 12 | QTFIS_F | 5'-CGATATCACCAGTACGCAAACTTTTATCAGCAGCACCGCCG-3' |
| 13 | QTIS_F | 5'-GATATCACCAGTACGCAAACTATCAGCAGCACCGCCGTGC-3' |
| 14 | Loopswap_F | 5'-GGCTTCGATATCACCAGTACGCCGCGCACCGAAGGCATCAGCAGCAC-3' |
| 15 | T7 Terminator | 5'-GCTAGTTATTGCTCAGCGG-3' |

**Table S11: List of strains and plasmids used in this study.**

| Name | Genotype | Source |
|---|---|---|
| **Strains:** | | |
| *E. coli K-12* MG1655 | Wild-type parent or wild type parent with a scar after homologous recombination | Michi Taga, University of California Berkeley |
| *E. coli* BL21 (DE3) | Overexpression strain | Thomas Pucadyil, IISER Pune |
| *Ec*FADS-Kan (WT-Kan) | *Ec*FADS in MG1655 genome with Kanamycin resistance cassette after homologous recombination | This study |
| *Loopswap* -Kan (LS-Kan) | *Loopswap Ec*FADS in MG1655 genome with Kanamycin resistance cassette after homologous recombination | This study |
| *E. coli* MG1655 K-12 *Loopswap 155-163* (*Loopswap)* | *Loopswap Ec*FADS mutant in MG1655 genome with a scar after homologous recombination | This study |
| **Plasmids:** | | |
| pET28a(+) | Empty vector | Michi Taga, University of California Berkeley |
| pLP001 | Wild-type *Ec*FADS on pET28a(+) | This study |
| pAS025 | Loopswap *Ec*FADS on pET28a(+) | This study |
| pAS026 | QTFCEGIS in the loop region in *Ec*FADS on pET28a(+) | This study |
| pAS027 | QTFCGIS in the loop region in *Ec*FADS on pET28a(+) | This study |
| pAS030 | QTFCIS in the loop region in *Ec*FADS on pET28a(+) | This study |
| pAS028 | QTCGIS in the loop region in *Ec*FADS on pET28a(+) | This study |
| pAS031 | QTFIS in the loop region in *Ec*FADS on pET28a(+) | This study |
| pAS029 | QTIS in the loop region in *Ec*FADS on pET28a(+) | This study |
| pAS001 | G23K-*Ec*FADS on pET28a(+) | This study |
| pAS002 | S165K-*Ec*FADS on pET28a(+) | This study |
| pAS003 | G23S-*Ec*FADS on pET28a(+) | This study |
